# Benchmarking Microbiome Methods for Improved Metagenomic Sequencing of Lower Respiratory Tract Samples

**DOI:** 10.64898/2026.10.16.759757

**Authors:** Jack T. Sumner, Erica M. Hartmann

## Abstract

Bronchoalveolar lavage is a common diagnostic strategy used to sample immune cells and microbiota of the lower respiratory tract. Metagenomic sequencing of raw bronchoalveolar lavage fluid reveals a low biomass microbial community often obscured by abundant host DNA. Here, we set out to determine an optimal strategy to robustly purify microbial DNA in excess of human DNA using a combinatorial lysis, host-depletion, DNA extraction approach. We identify strategies that improve metagenomic sequencing success that are commensurate with purification challenges that reduce signal from respiratory microbiota. Finally, we present a general framework to further improve microbial sequencing of lower respiratory tract samples in a robust and reproducible manner.

## 1.2 BACKGROUND

Cost-effective high throughput sequencing technology has led to unprecedented advances in microbial ecology.^1,2^ Culture-independent methods like sequencing have highlighted prevalent yet uncultivable microbiota that traditional methods fail to identify. Targeted sequencing approaches, including 16S rRNA gene sequencing, are a cost-effective means to profile bacterial communities and have thus been implemented in a wide range of environments^3,4^. However, limited taxonomic resolution and lack of phenotypic insights are major deficits to targeted sequencing approaches in comparison to culture-dependent methods.^5,6^

Shotgun metagenomic sequencing captures mixed DNA from biological community samples.^7^ Unlike targeted approaches, shotgun metagenomics is not limited to a single locus and can ideally resolve entire genomes for downstream analysis. Analysis of encoded genes and their annotations can be used to infer the functional potential of complex microbial communities. Although shotgun metagenomics has been implemented in many sample types,^8–10^ certain sample characteristics limit the efficiency of this method to resolve genomic information.^11,12^

Bronchoalveolar lavage is a commonly-used diagnostic tool that is capitalized on to harvest lower respiratory tract microbiota.^13^ Metagenomic sequencing of bronchoalveolar lavage fluid can be used both as a diagnostic to identify unknown pathogens^14^ and as a tool to study the lung microbiome in disease.^15,16^ Attempts to use shotgun metagenomic sequencing on raw bronchoalveolar lavage fluid result in poor microbial resolution and sequencing efficiency.^11,17–21^ Bronchoalveolar lavage yields patient-derived DNA at much greater concentrations than microbiome-derived DNA, likely due to the low biomass nature of these microbial communities. Thus, novel sample processing pipelines that deplete patient-derived DNA while retaining microbiome-derived DNA must be developed to improve sequencing success and to further unveil microbial function in disease.

In this study, we propose a combinatorial optimization framework strategy of the entire sample processing workflow to robustly improve metagenomic resolution via host-depletion while mitigating bias. By examining the impact of orthogonal methods at each step of the workflow, we present a near comprehensive validation scheme that may be translated to other human biopsies. We present specific data to showcase the benefits and limitations of host-depletion techniques on improving metagenome sequencing success, and how the interpretation of success may be influenced by artifacts occurring at different stages of the workflow.

## 1.3 RESULTS

### SAMPLE COLLECTION & TREATMENT

#### Sampling Overview

Sampling of the lower respiratory tract was performed using bronchoalveolar lavage (BAL) by appropriate healthcare providers (e.g., pulmonologists) as part of routine clinical care. BAL is a relatively minimally invasive sampling technique that involves instillation of saline (0.9 N NaCl) into the alveolar space and subsequent aspiration of BAL fluid (BALF) through either a specialized catheter or a fiberoptic bronchoscope for diagnostic purposes. BALF samples were collected from patients with suspected pneumonia in the intensive care unit at Northwestern Memorial Hospital. After appropriate clinical use, residual material from eight BALF samples was collected for methods optimization research.

### PRE-LYSIS TREATMENT IMPACTS NON-HOST METAGENOMIC SEQUENCING DEPTH

During collection, aliquots of the 8 BALF samples were treated in a combinatorial two-step manner (Figure 3). Primary treatment processing included centrifugation of BALF sample aliquots followed by either resuspension of the pellet (“resuspend”; [Re]) or discarding of the supernatant and resuspension of the pellet in nuclease-free water to promote osmolysis of host cells (“osmolysis”; [Ly]). A no-treatment control was set on ice for downstream processing (“native”; [Na]). Secondary treatment processing was then performed combinatorially on each of the primary treated sample aliquots. BALF sample aliquot received treatment with either DNaseI or PMA to deplete accessible DNA. An additional no-treatment control group was included at this step (i.e., native). This yielded 9 aliquots for each sample, each with a different combination of the three primary and three secondary treatment processing groups. Downstream DNA extraction was performed in seven randomized batches of up to 11 BALF sample aliquots for a total of 72 BALF aliquot DNA samples and 7 blank extraction controls. A no-template (i.e., water) and a standard of known microbial DNA content (i.e., ZymoBIOMICs Microbial Community DNA Standard) were included during sequencing library construction as negative and positive control for library construction.

**Figure 1.**
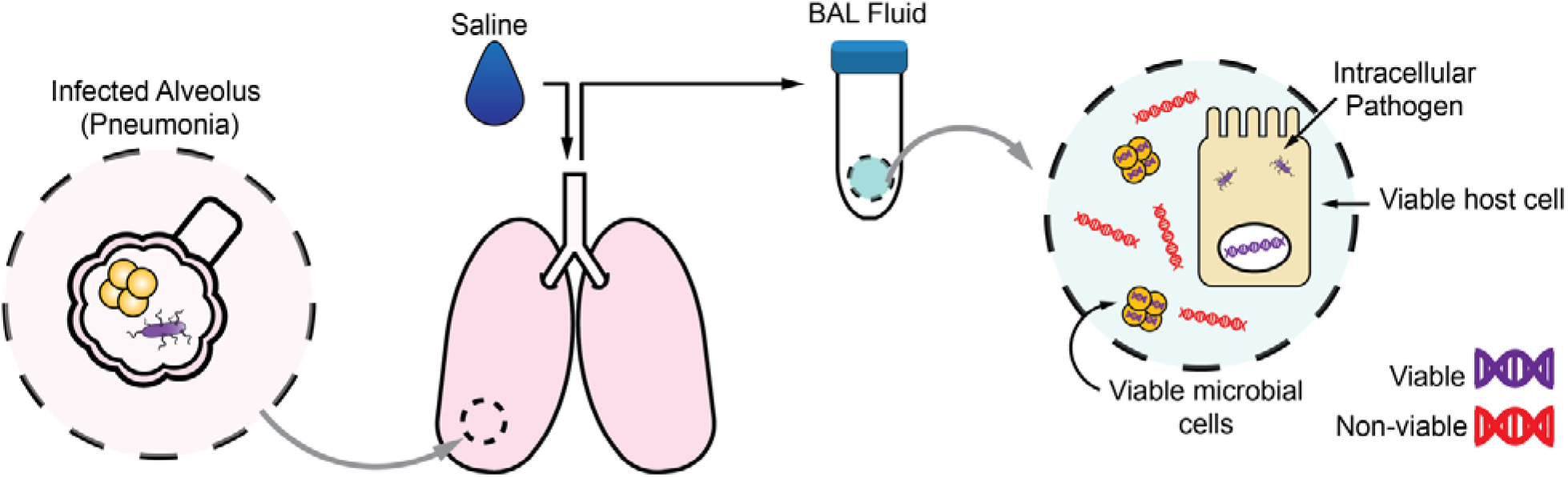
Illustration of bronchoalveolar lavage fluid. Saline solution (90% NaCl) is instilled into the lungs of patients with pneumonia for diagnostic testing. Fluid is then aspirated and used for downstream clinical testing and research applications. Bronchoalveolar lavage is a complex matrix of material from mucosal lining that retains free and intracellular DNA from patient-cells and microbiota.

**Figure 2.**
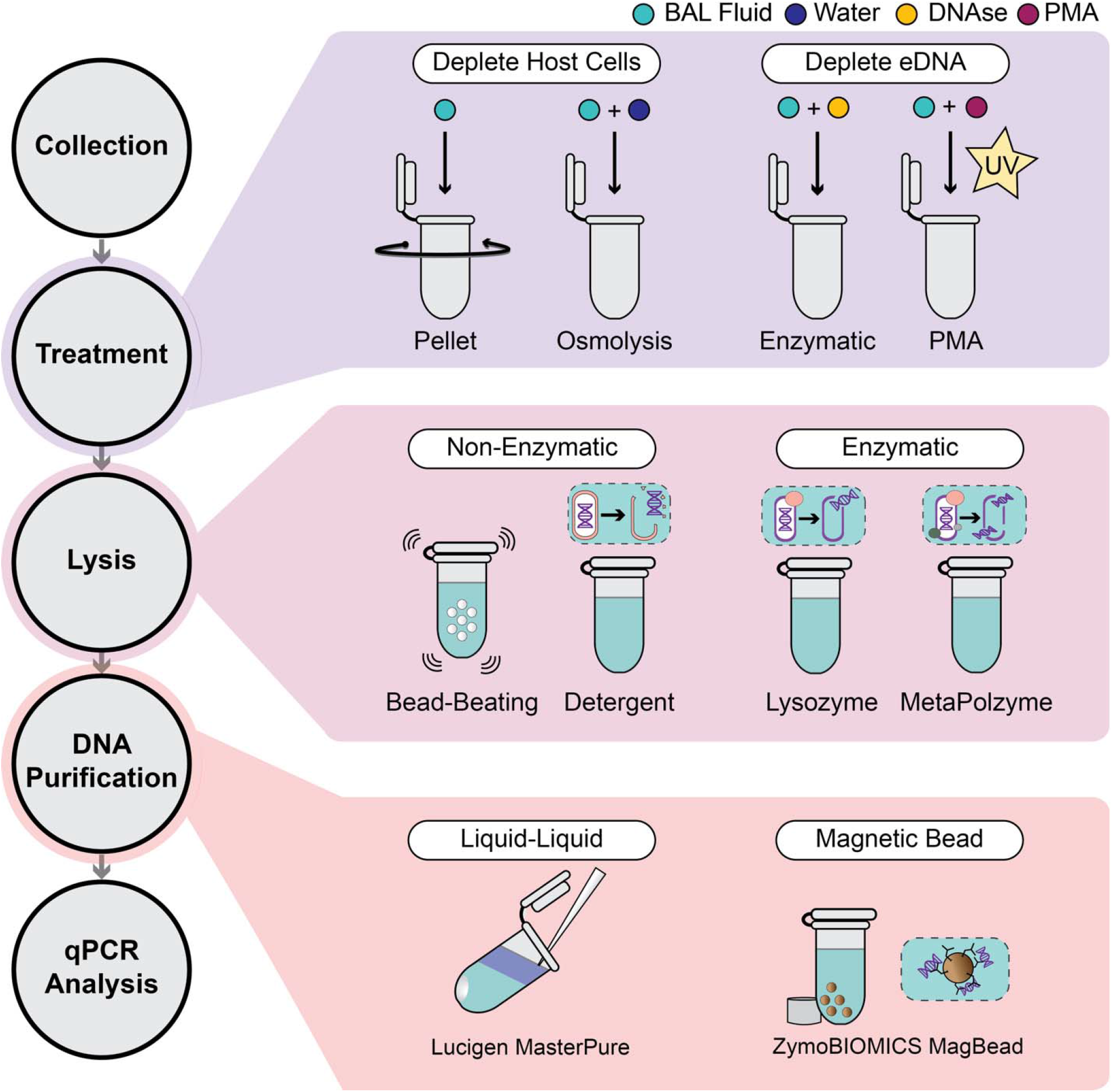
Schematic of treatments to improve microbial coverage in bronchoalveolar lavage fluid metagenomes. Selective treatment strategies can be applied to target host cells or extracellular DNA for patient-DNA depletion. Lysis methods have reported biases and efficiency; screening different approaches will identify potential pitfalls in microbiota yield. DNA purification has tradeoffs in yield and throughput such that automated approaches may not yet be suitable for low-biomass microbial communities.

**Figure 3.**
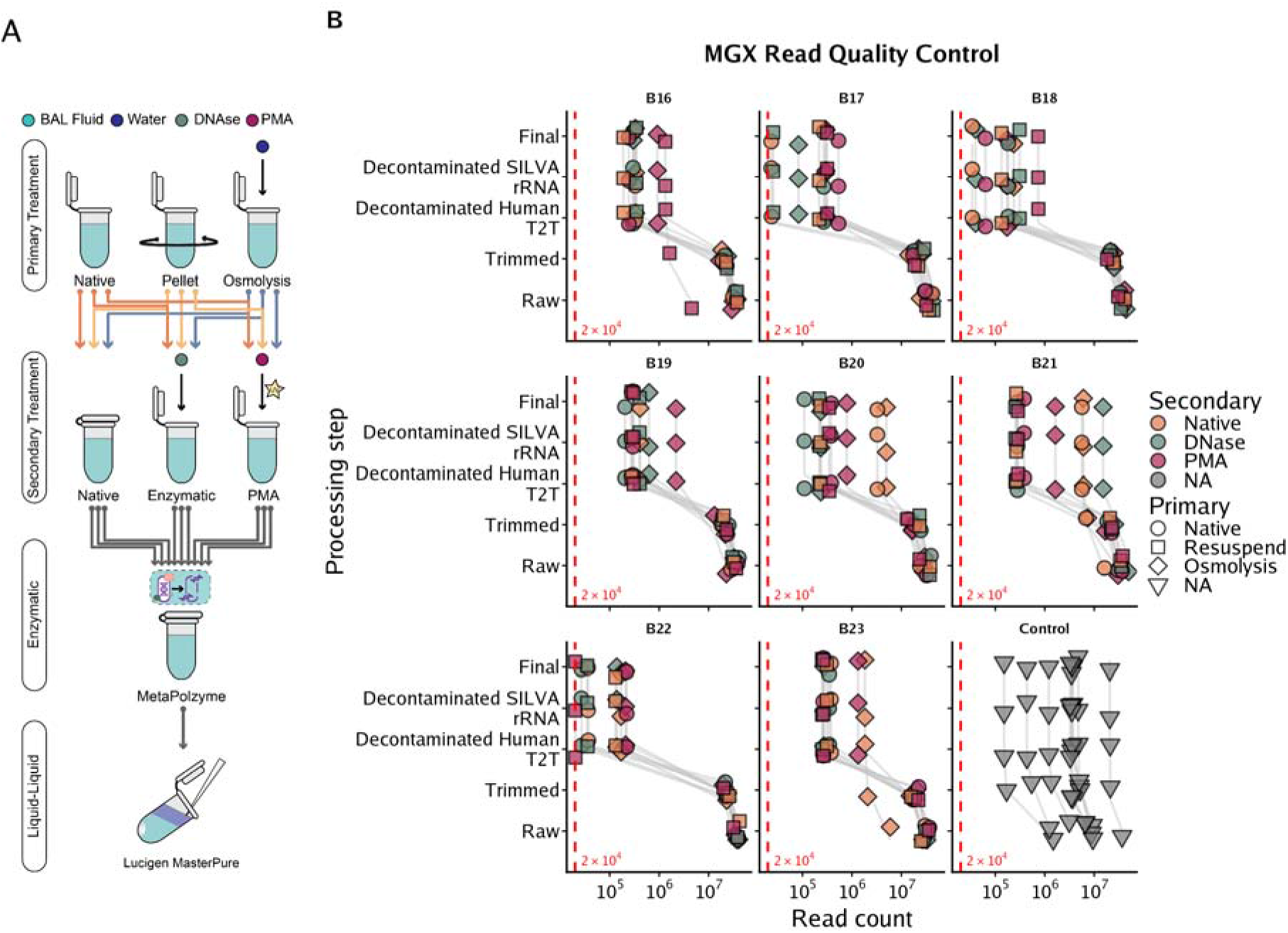
BALF samples aliquots were processed using different pre-lysis conditions that variably impacted sequencing depth. (**A**) Overview of sample processing pipeline used to generate the 9 pre-lysis treatment combinations for each of the eight BALF samples. DNA from extracted from BALF sample aliquots were subsequently used in quantitative PCR and sequencing assays. (**B**) Visualization of metagenomic reads removed at each quality control step. Quality control was performed using KneadData. Within KneadData, reads were trimmed (“Trimmed”) followed by serially aligning to the human genome (“Decontaminated Human T2T”) and SILVA rRNA gene database (“Decontaminated SILVA rRNA”). Control samples included blank extraction controls as well as positive (ZymoBIOMICS microbial community DNA standard) and negative (water) controls for library construction; control read are shown in the lower right-hand corner. The control sample with the highest and lowest number of raw reads corresponds to negative and positive library construction control, respectively. (Acronyms: BALF = bronchoalveolar lavage fluid; PMA = propidium monoazide).

Quality control for metagenomic sequencing data was performed using KneadData (Figure 3B, Supplemental Table 1). Briefly, reads were trimmed of sequencing adapters, low-quality bases, and short reads using Trimmomatic. Reads were then serially aligned to the human reference genome and a database of rRNA genes to deplete host reads and reads irrelevant to marker gene analysis. The number of reads surviving each step of the analysis are visualized in Figure3B. The results indicate that BALF samples processed with primary osmolysis treatment or secondary DNase and PMA tended to have a smaller fraction of human-derived reads than other processing groups. This indicates that host-depletion strategies are likely reducing the fraction of host-associated reads from the pool of sequenced DNA fragments.

In some cases, the final set of quality-controlled reads failed duplication assessment in FastQC in a way that is relevant to interpretation of host-depletion efficacy. From these samples, we observe duplication levels greater than 50% from FastQC outputs. These samples include aliquots from BALF sample B20 (i.e., NaNa, LyNa), B21 (i.e., NaNa, LyNa, LyDNase) and B23 (i.e., LyNa). Re-inspection of the TapeStation traces from prepared libraries of these samples indicated the presence of a secondary, spiked peak at ∼150 bp in addition to the larger broad peak typical of metagenomic libraries. A peak of this size and amplitude is often indicative of primer dimers. This suggests that, in these samples, artifacts from the sequencing library construction are likely contributing to the fraction of non-host reads, thereby artificially enhancing host-depletion efficacy in these samples. Thus, low-levels of primer-dimers (or similar artifacts) that would be acceptable for metagenomic sequencing of high-microbial biomass samples (e.g., stool) may hinder sequencing success in samples with high host-load and low-microbial biomass. Furthermore, the presence of low-level primer dimers may be conflated for improved microbial sequencing depth and host-depletion treatment effects unless specifically taken into consideration.

### PRE-LYSIS TREATMENT IMPACTS HUMAN AND BACTERIAL BIOMASS IN METAGENOMES

Bacterial and human load were determined using quantitative polymerase chain reaction (qPCR) targeting the 16S rRNA and PTGER2 genes, respectively. Copies of 16S rRNA gene were slightly lower in water-incubated samples than in native samples (FDR *P* < 0.05; Figure 4A); PMA treated sample retained lower concentrations of 16S rRNA gene copies than did DNase treated samples (FDR *P* < 0.001; Figure 4B). A similar trend was observed for PTGER2 gene concentration, a marker for the human genome (Figure 4F-G). Osmolysis treated samples and PMA treated samples were significantly depleted of the PTGER2 gene relative to native treatment and primary resuspended or secondary DNase treatment, respectively (FDR *P* < 0.001; Figure 4F-G). Concordantly, human reads represented a slightly smaller fraction of osmolysis treated samples (Figure 4K; FDR *P* < 0.05). Human reads represented a significantly smaller fraction of the sequenced metagenome in samples treated with PMA relative to DNase treated samples (Figure 4L; FDR *P* < 0.05). These data indicate that treatment with PMA decreases the yield of human DNA and, to a much lesser extent, of bacterial DNA in BALF samples.

**Figure 4.**
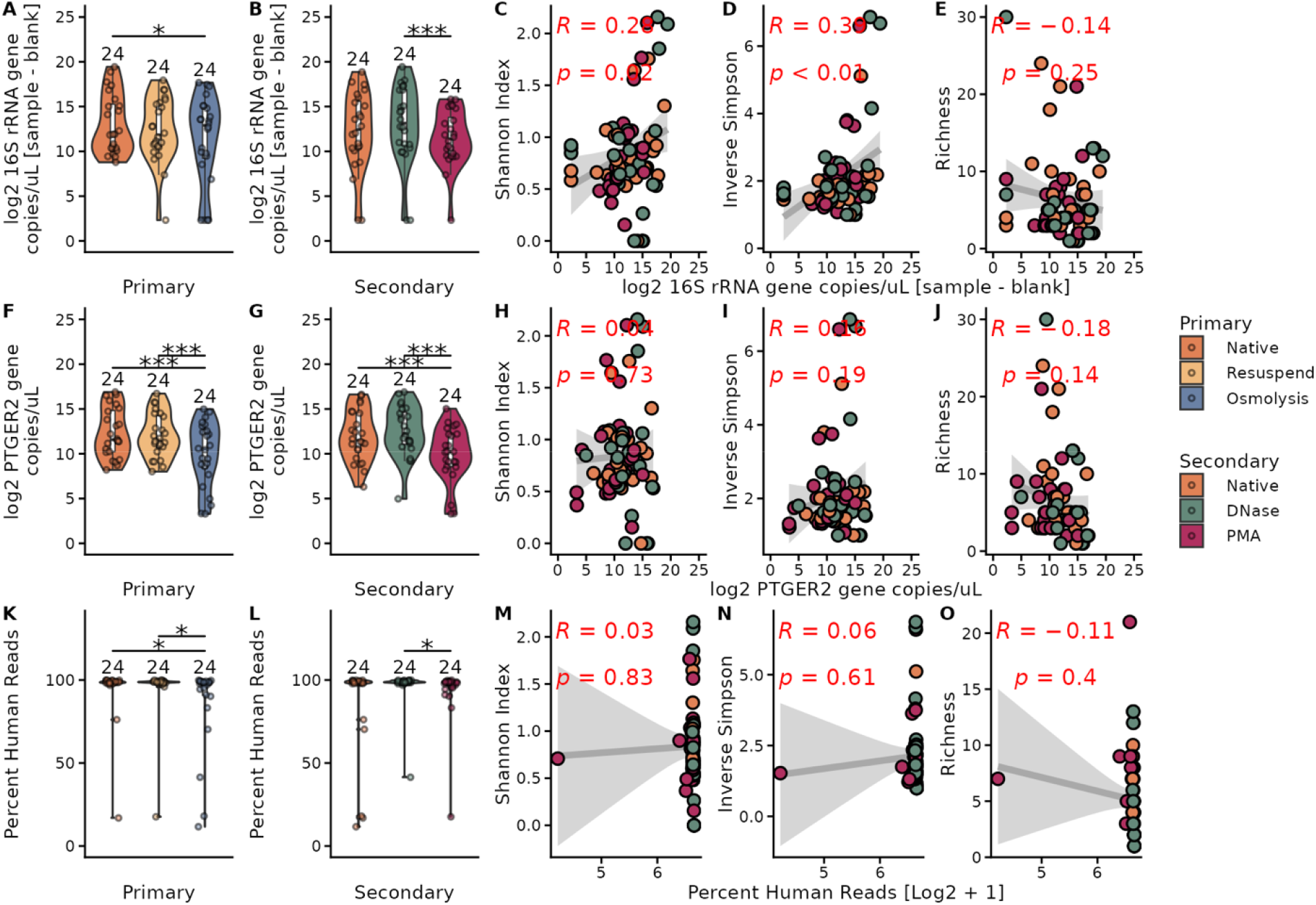
Effects of pre-lysis treatment on bacterial and human biomass in BALF samples. (A-B) Effects of (**A**) cell-targeted primary treatment and (**B**) DNA-targeted secondary treatment on 16S rRNA gene copies recovered from BAL samples. Copy numbers are adjusted by batch-specific extraction control measurements. (C-E) Associations of recovered 16S rRNA gene copies with metrics of alpha diversity, including (**C**) Shannon diversity, (**D**) inverse Simpson, and (**E**) species richness. (F-G) Effects of (F) cell-targeted primary treatment and (G) DNA-targeted secondary treatment on PTGER2 gene copies recovered from BAL samples. (H-J) Associations of recovered PTGER2 gene copies with metrics of alpha diversity, including (**H**) Shannon diversity, (**I**) inverse Simpson, and (**J**) species richness. (K-L) Effects of (**K**) cell-targeted primary treatment and (**L**) DNA-targeted secondary treatment on human read fraction in shotgun metagenomic libraries generated from BALF samples. (M-O) Associations of log2 transformed human read fraction with metrics of alpha diversity, including (**M**) Shannon diversity, (**N**) inverse Simpson, and (**O**) species richness. (C-E, H-J, M-O) Linear regressions visualized as grey line with shaded region as 95% confidence interval; red text represents results of regression analysis. Boxplots stars in violin plots (A-B, F-G, K-L) are the results of non-parametric Wilcoxon Signed-Rank tests (paired, A-B, F-G, K-L). (**Boxplot configuration**: Center line = median, box limits = upper and lower quartiles, whiskers = 1.5x interquartile range. **Stars**: * = FDR *P* < 0.05, ** = FDR *P* < 0.01, *** = FDR *P* < 0.001. **Acronyms**: AST = arcsine square root transformation).

Several alpha diversity metrics were evaluated in conjunction with qPCR and metagenomic sequencing depth data. The inverse Shannon index and Simpson index positively associated with 16S rRNA gene concentration (range *P* < 0.05-0.01; Figure 4C-D); no association with species richness was detected (Figure 3E). Neither Shannon (Figure 4H), inverse Simpson (Figure 3I), nor species richness (Figure 4J) associated with PTGER2 gene copy concentration (*P* > 0.05). No associations were detected between the human fraction of metagenomic reads and the observed alpha diversity (Figure 4M-O). Together these data indicate that increasing bacterial biomass yield in metagenomic DNA extraction, and the resultant depletion of human-derived reads, was moderately associated with detection of more diverse respiratory tract microbiota.

### CONTAMINANT ABUNDANCE VARIES IN A SAMPLE AND TREATMENT-DEPENDENT MANNER

Contamination was abundantly detected throughout the studied samples (Figure 5). The prevalence methodology from the Decontam R package was used to identify contaminant species based on the taxonomic profiles found in negative controls (see Methods for details). Additional contaminants were added based on visual inspection of the taxonomic profiles and prior knowledge of the lower respiratory tract microbial community. These contaminants include other species of the genus *Lysobacter*, for which the close relative *Lysobacter enzymogenes* was identified as putative contaminant with high relative abundance. Additionally, the species *Methylibium petroleiphilum* was manually included as a contaminant. Although *L. enzymogenes* was typically the highest relative abundance among blank extractions for the first six batches (∼45-80% relative abundance), *M. petroleiphilum* was the greatest contributor to the seventh blank extraction control (∼75%) and no-template library construction control (∼35%). Furthermore, *M. petroleiphilum* was detected at relatively low levels in the first six batches (∼10^-1^ relative abundance). This suggests that contamination differs by batch and may correspond to differences in when the contaminant was introduced (extraction vs library construction) and differences in reagent batch contamination (e.g., MetaPolzyme lot). This species was only identified as a contaminant upon increasing the Decontam threshold to the maximum recommended value of 0.5, which also led to identification of likely false positive contaminants. The human microbiota of *Staphylococcus pasteuri* and the uncultured Lawsonellaceae member (SGB3663) were excluded from the contaminant list as they were of consistent low abundance among blank extractions and are potentially derived from BALF samples. Despite this moderately conservative approach, contamination was consistently detected across BALF samples.

**Figure 5.**
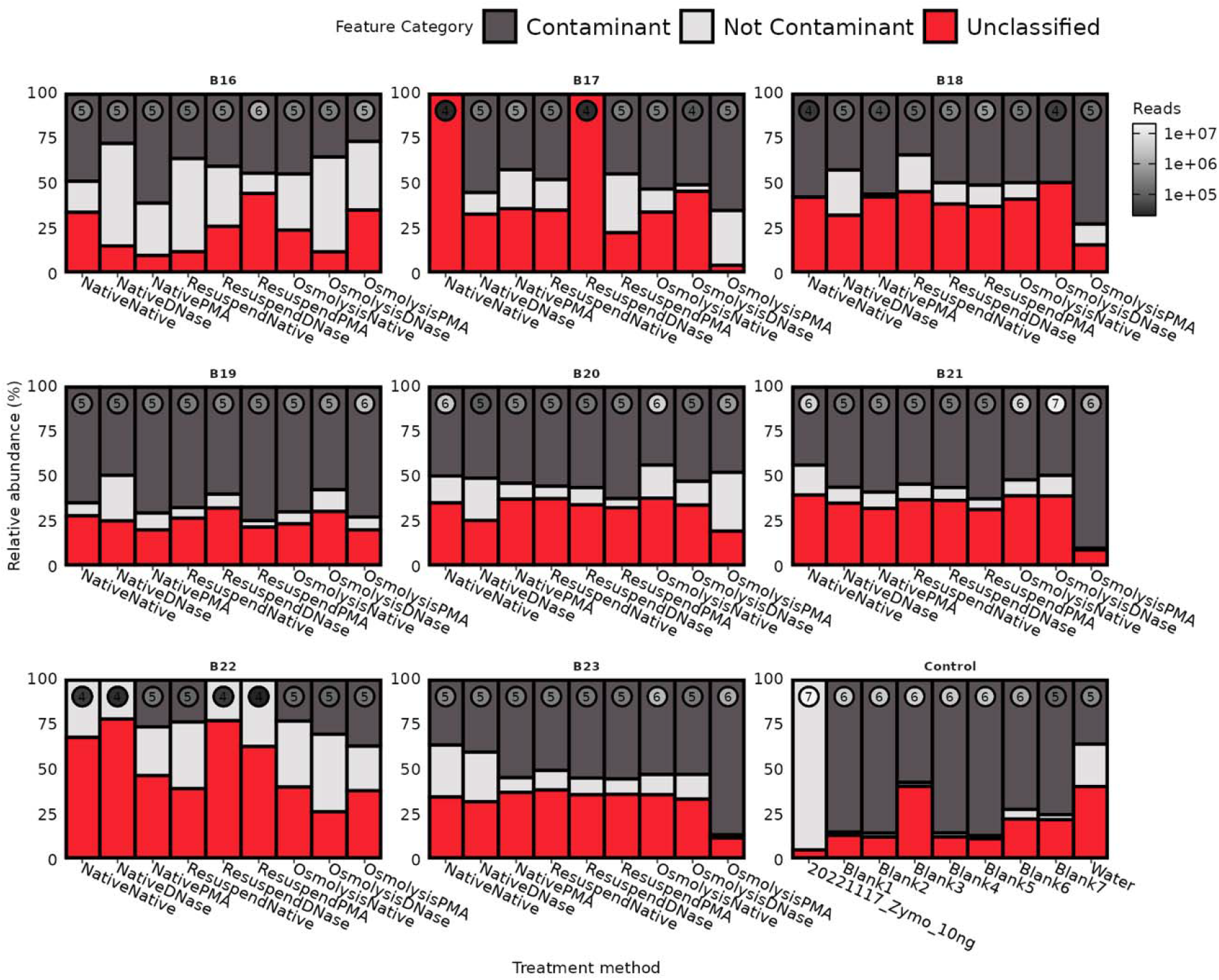
Putative contaminant abundances vary by BALF sample and pre-lysis aliquot treatment. Stacked barplots visualizing the relative abundance of putative contaminants identified using the Decontam R package and manual curation (see Methods for details). Points towards the top of each sample are colored by the final number of quality-controlled reads. For ease of interpretation, the order of magnitude of total reads (e.g., log10 floor rounded to the nearest whole number) is displayed as text within each point.

Overall, metagenome contamination tended to differ between BALF samples. This is consistent with the previous observation that bacterial biomass and microbial profiles are highly distinctive between individual subjects. So, samples of lower true microbial biomass would likely be more susceptible to contamination as the pool of available microbial DNA for sequencing would be lower than in samples of higher microbial biomass. In most cases (n = 6/8 samples), treatment with osmolysis followed by PMA depletion increased the relative fraction of contaminants compared to no treatment (Figure 5). Sample aliquots with few surviving reads post host depletion (i.e., on the order of 10^4^ reads) had no discernable contamination, although were mostly composed of unclassified reads (n = 6 aliquots; Figure 5). This is notable as MetaPhlAn4 typically requires at least this order of magnitude of reads to identify species. The fraction of non-contaminant taxa that were classified varied across sample aliquots, sometimes relatively increasing in treated samples compared to native controls. These data suggest that DNA-depletion methodologies often increase susceptibility to contamination, e.g., from reagents, while simultaneously altering the fraction of sequenced material from true lower-respiratory tract microbiota. This led us to hypothesize that variation in non-contaminant microbiota detection was associated with treatment groups.

### PRIMARY AND SECONDARY TREATMENT CONTRIBUTE TO VARIATION IN JACCARD SPACE

To determine the level of variation between aliquots processed with different pre-lysis treatment strategies, we implemented a distance metric-based approach. The benefit of this approach, over relying on differences in specific taxa, is that we can more easily adjust to differences from baseline (raw or native) samples in a low-dimensional space. This is key for BALF samples because taxa are typically not expected to be highly conserved between samples from different subjects, contrary to what one might expect in other body sites (e.g., the gut). Therefore, to detect taxa that are differentially abundant with treatment effects, (1) random effects models implementing subject-specific identifiers and a (2) much greater sampling depth, both technical and biological replicates, to account for this would be required. However, variation in pairwise dissimilarity was associated with specific treatment group comparisons.

BALF sample aliquots were compared to other aliquots (derived from the same sample) receiving the same primary treatment (Figure 6A) and to those receiving a different primary treatment (Figure 6B). Within-treatment dissimilarity was similar across primary treatments, with a small difference detected between within-osmolysis and within-resuspended treatments (FDR *P* < 0.05; Figure 6A). Pairwise Jaccard dissimilarity was lower between native-resuspend and resuspend-osmolysis comparisons than it was between other comparisons (FDR *P* < 0.05; Figure 6A). This indicates that native and osmolysis treated aliquots were more dissimilar to each other relative to the other between-group primary treatment comparisons.

**Figure 6.**
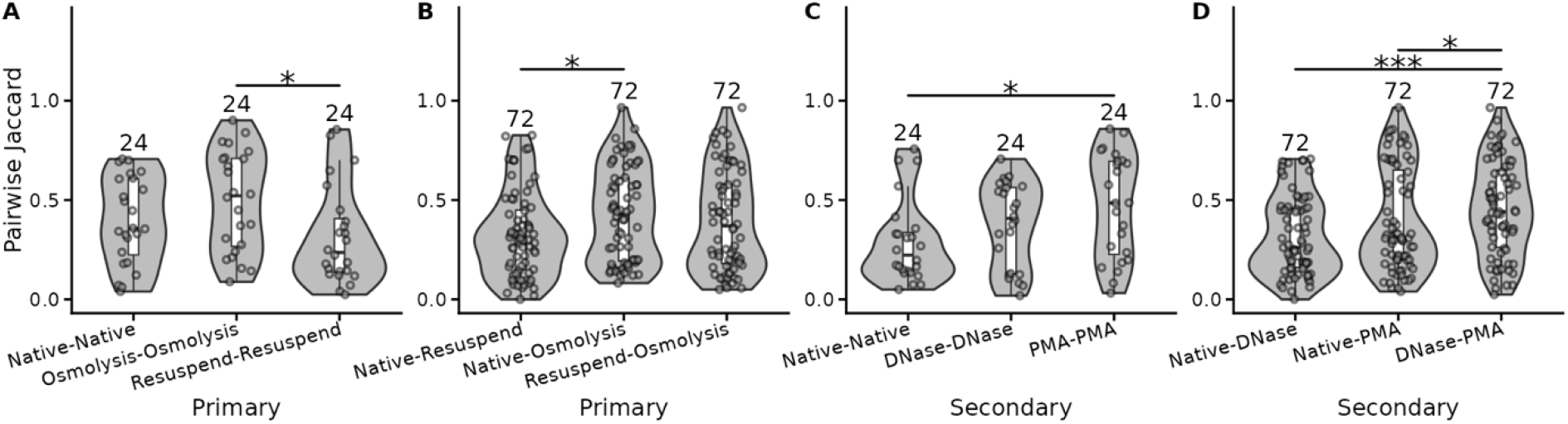
Pairwise distance comparisons between treatment groups. Pairwise Jaccard distance between BALF sample aliquots (within individual) (**A**) within the same primary treatment group and (**B**) between different primary treatment groups. Pairwise Jaccard distance between BALF sample aliquots (**C**) within the same secondary treatment group and (**D**) between different secondary treatment groups. Comparisons are limited to aliquots from the same BALF sample (e.g., B16 NaNa is compared to B16 NaPMA and not B17 NaNa in leftmost violin of A). Boxplots stars in violin plots are the results of non-parametric Wilcoxon Signed-Rank (paired) test. (**Boxplot configuration**: Center line = median, box limit = upper and lower quartiles, whiskers = 1.5x interquartile range, points = outliers. **Stars**: * = FDR *P* < 0.05, ** = FDR *P* < 0.01, *** = FDR *P* < 0.001).

Secondary treatment followed a similar pattern of associations. Within-treatment dissimilarity was highest in PMA treated samples compared to natively treated samples (FDR *P* < 0.05), but otherwise consistent between remaining secondary treatment comparisons (Figure 6C). Dissimilarity was greater between DNase and PMA treated samples than it was between native and DNase treated or native and PMA treated samples (FDR *P* < 0.001, FDR *P* < 0.05; Figure 6D). This indicates that DNase and PMA treated aliquots were more dissimilar to each other than were native and DNase treated aliquots. It suggests that DNase and PMA treatment may lead to recovery of different facets of the microbial community.

### SPECIES-LEVEL METAGENOMIC PROFILES ARE ALTERED BY PRE-LYSIS TREATMENT

BALF sample taxonomic profiles differ in response to primary (cell-targeted) and secondary (DNA-targeted) pre-lysis treatments. Overall, the microbiota signatures of BALF aliquots from the same sample tended to resemble each other more than aliquots from other samples (Figure 7A). For instance, metagenomic sequencing results indicated that all nine aliquots of BALF sample B22 were positive for *Metamycoplasma hominis*, which was observed only sporadically in three other BALF samples (i.e., B16 native-PMA and osmolysis-PMA; B19 native-DNase; and B23 osmolysis-PMA). In BALF sample B16, aliquots tended to be positive for at least one species of the genus *Lactobacillus* and *Limosilactobacillus* (Figure 7A; n=8/9 for B16 aliquots both genera). Aliquots from this sample displayed greater specie richness than other BALF samples, independent of the treatment group. Together, this indicates that sample-specific microbial signatures represent a great source of variation in BALF samples, as previousl described.^16^

**Figure 7.**
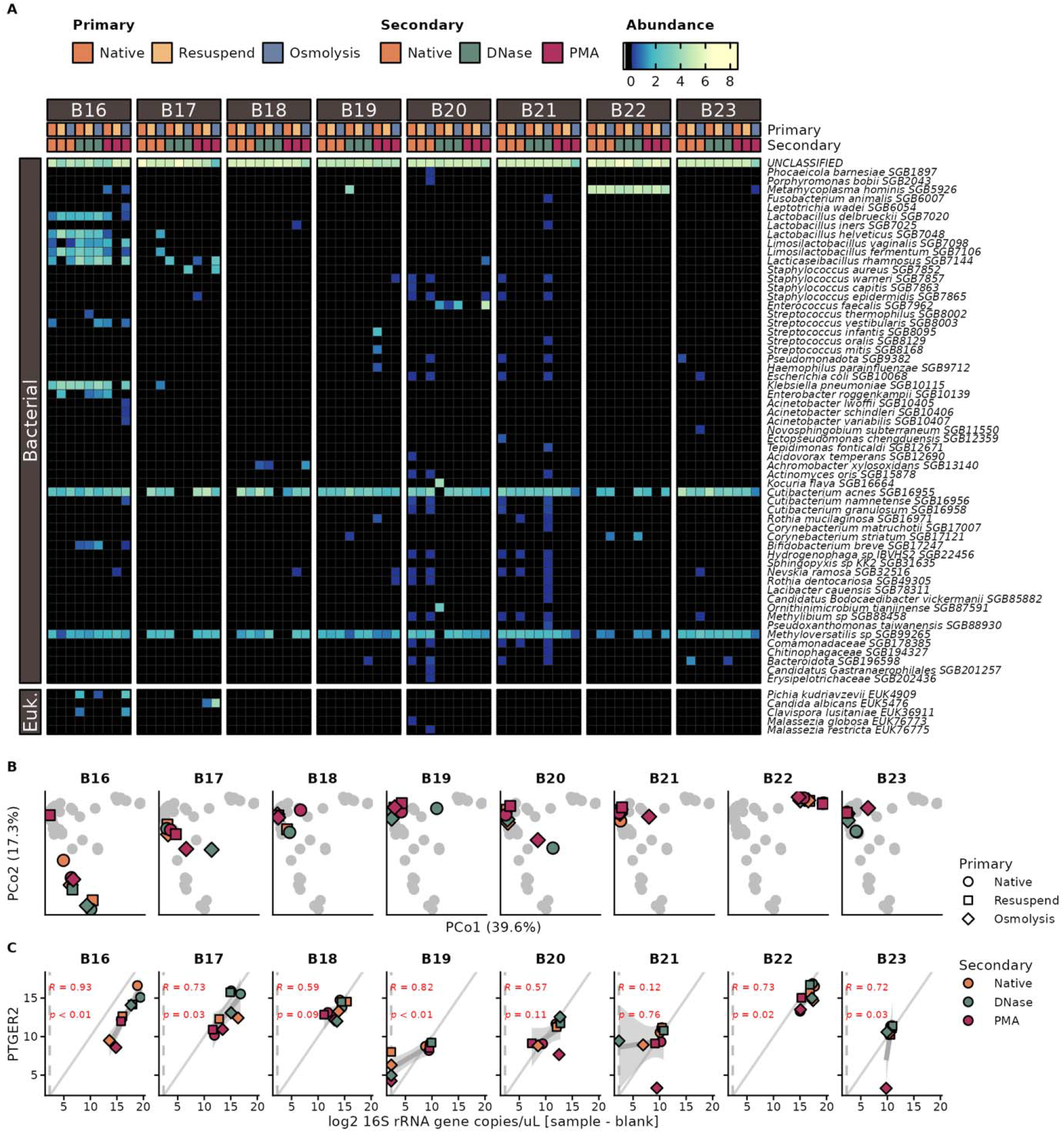
Impact of pre-lysis treatment on BALF taxonomic profiles at sample-level resolution. (**A**) Heatmap visualizing species-level taxonomic profiles generated by MetaPhlAn4. Grouped columns are BALF samples labelled with their sample index in the grey boxes; columns are aliquots of BALF samples. Aliquot treatment is as depicted in the top annotation rows (i.e., Primary, Secondary). Features (rows) are grouped by taxonomic domain for ease of interpretation. Feature order is the result of hierarchical clustering using Bray-Curtis dissimilarity as the distance metric (dendrogram is omitted). (**B**) Principal coordinate analysis of taxonomic profiles using the Bray-Curtis dissimilarity. Shape indicates cell-targeting primary treatment, and color is indicative of DNA-targeting secondary treatment. Gray dots in the background are the shadows of points from all subpanels combined. (**C**) Regression analysis of 16S rRNA gene and PTGER2 gene copy numbers from extracted metagenomes. Panels represent distinct samples. The solid grey line is a slope of 1 to showcase balance between 16S rRNA (bacterial) and PTGER2 (human) genes. The dashed grey line indicates the limit of quantification for 16S rRNA gene qPCR assay. Linear regressions visualized as dark grey line with shaded region as 95% confidence interval; red text represents results of regression analysis. (Acronyms: Euk. = Eukaryotic)

Treatment of BALF aliquots tended to increase metagenomic detection of microbiota with clinical interest. In 2 BALF samples, treatment had minimal discernable impact on microbial taxonomic profiles (Figure 7A; samples B18 and B23). In all other BALF samples, at least one treatment group increased specie detection. Treatment led to increased detection of pathogenic microbiota in half of the BALF samples (Figure 7A). This effect was attributed to osmolysis followed by PMA (LyPMA) treatment in three BALF samples. In BALF sample B16, LyPMA treatment and native-DNase treatment (NaPMA) promoted detection of the pathogenic yeast *Clavispora* and *Pichia*, the latter of which was also detected by LyDNase treatment in the same BALF sample. Furthermore, LyPMA treatment promoted detection of *Candida albicans* and *Staphylococcus aureus* in sample B17. LyPMA and DNase treatment promoted detection of *Enterococcus faecalis* in sample B20; relative abundance of *E. faecalis* was notably higher in LyPMA than in DNase treated samples for this BALF sample (Figure 7A). Application of DNase without primary treatment led to the detection of *Metamycoplasma hominis* in one sample (B19). *Enterobacter roggenkampii*, an opportunistic pathogen, was detected in four aliquots of sample B16 (i.e., ReNa, ReDNase, LyDNase, NaPMA). Notably, the pathogen *Klebsiella pneumoniae* was detected independent of processing treatment for aliquot B16 (n = 8/9 aliquots). The complete sample-level taxonomic profile is depicted in Figure 7A. Overall, response to treatment was heterogenous across BALF samples, but application of secondary treatment led to increased microbiota detection in metagenomic sequencing data.

Principal coordinate analysis was used to visualize variation between samples and treatment aliquots (Figure 7B). Treatments that increased pathogen detection were often farther from other aliquots of the same sample in principal coordinate space (e.g., B19 NaDNase). Therefore, the results of principal coordinate analysis are concordant with the shifts in species detection described in the previous paragraph. This indicates that host depletion in BALF samples alters microbiota signatures in metagenomic sequencing data.

The degree to which treatment alters the microbiota profile is contextualized by the efficacy and specificity of host depletion. To that end, we performed linear regression analysis against PTGER2 and 16S rRNA gene copy concentrations data (Figure 7C). We expected effective host depletion to decrease PTGER2 gene copy concentration, while specific host depletion would have minimal effect on 16S rRNA gene copy concentration. Therefore, the ratio of bacterial to host load in metagenomic DNA extractions would increase and lead to increased metagenomic sequencing success. LyPMA tended to have high efficacy and specificity host depletion based on qPCR data. This is most evident in samples B16, B20, B21, and B23 as well as in samples B17 and B22 to a lesser degree. This does not always translate to improved sequencing success in a consistent manner, as with sample B23, which had the best response to host-depletion in qPCR data but minimal response in the sequencing profiles. Conversely, PTGER2 gene copy concentration (but not 16S rRNA gene copy concentration) was approximately halved in the B17 LyPMA treated aliquot relative to untreated controls. It was also this sample aliquot that had improved detection of *S. aureus*. The resulting data indicate that qPCR-based screening may be used to measure host-depletion efficiency and specificity as a means of predicting metagenomic sequencing success.

### SPORADICALLY DETECTED FEATURES ARE OF RELATIVELY LOW ABUNDANCE

We hypothesized that features sporadically detected across treatment groups would be of low relative abundance. To test this hypothesis, we first examined the relationship between species prevalence and its maximum abundance within a BALF sample and across all samples (Figure 8A). The relationship between species prevalence and its mean abundance was also assessed (Figure 8B). It is clear from these regressions that species with a lower within-sample prevalence tended to have lower mean abundance, even when a feature is detected at high abundance.

**Figure 8.**
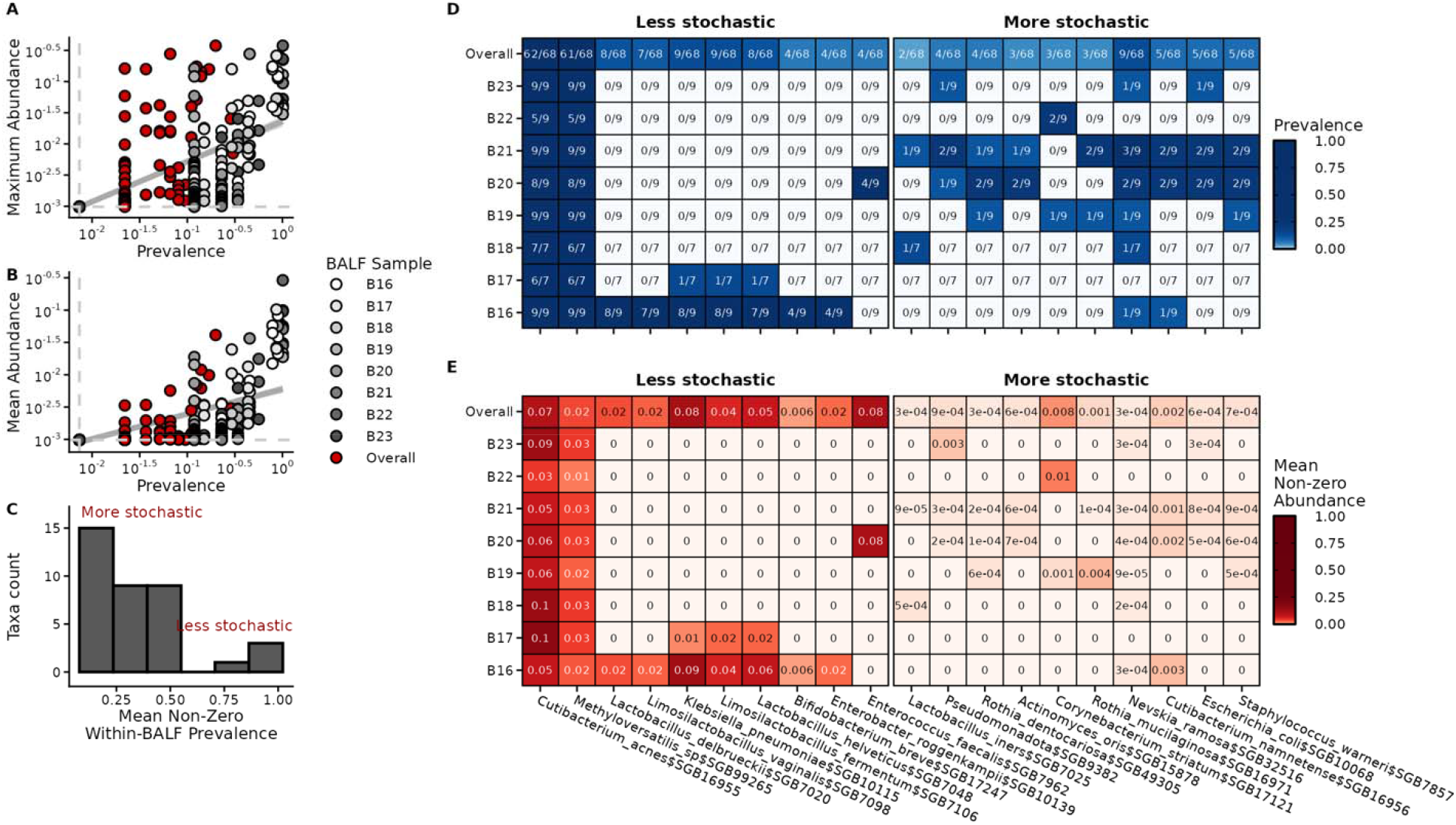
Stochastic feature detection tends to be associated with lower-level abundance than features that are not stochastically associated. (**A**) Relationship between species prevalence and its maximum relative abundance. Grey points are within-BALF sample values (e.g., species prevalence in B16 sample aliquots) while red points are across all samples and aliquots (“overall”). Dotted lines represent ½ the lowest non-zero prevalence and abundance values; zero-values were imputed to this number to keep all points on the graph. (**B**) Relationship between species prevalence and its mean relative abundance. (**C**) Mean within-BALF feature prevalence of detected features (MWBP; i.e., prevalence values greater than zero in each BALF). Here, a higher MWBP indicates the feature was more consistently detected across a set of BALF sample aliquots than features with a lower MWBP. Count is number of species in each histogram bin. Species detected in at least any two aliquots were selected for inclusion. (**D**) Prevalence and (**E**) mean non-zero abundance of the highest (“Less stochastic”) and lowest (“More stochastic”) species by MWBP. Mean non-zero abundance is shown here to avoid conflating low-abundance with low-prevalence.

To rank species by how sporadically they were detected, we calculated the mean within-BALF prevalence of values greater than zero (Figure 8C). We posit that the features with the greatest mean non-zero within BALF prevalence (MWBP) score tend to be more sporadically detected in our dataset. The top and lowest species by MWNP are shown in Figure 8D. Taxonomic features with higher MWNP score tended to have greater relative abundances compared when detected (∼10^-2^-10^-1^) compared to taxonomic features with lower MWNP score (∼10^-4^; Figure 8C). Notably, most features were of low relative abundance (i.e., less than ∼1%) independent of MWNP score; this was expected as profiles were not renormalized after contaminant removal.

To further test the hypothesis that detection of taxonomic features is a stochastic process, we evaluated the empirical distribution of within-BALF prevalence against a Poisson distribution. The Poisson distribution is commonly used to model stochastic processes in time. One common example of a Poisson process is transcription factor binding to a given cis-regulatory site encoded in the genome. Here, we fit the Poisson, gamma, and negative binomial distributions to scaled density of the within-BALF prevalence (Figure 9). In some instances, the Poisson distribution appeared to fit within-BALF prevalence well (e.g., B17, B20, B21, B22, somewhat B19). In other instances, the Poisson distribution was clearly a poor fit (e.g., B16, B18, B23, Controls). Notably, samples that poorly fit the Poisson distribution had a bi-modal empirical distribution of within-BALF species prevalence. We thus hypothesize that within-BALF prevalence distributions of these samples would be better explained by a mixture model combining a Poisson distribution and one other distribution at the low and high ends of within-BALF prevalence. The detection of bimodal distributions that at least partially do not fit a Poisson distribution is consistent with our analysis of MWNP score, which detected some taxa that were consistently detected at higher relative abundances than taxa of lower MWNP score. Overall, these data indicate that low-abundant microbiota are detected in a manner consistent with a Poisson process. This suggests that detection of rare taxa was likely due to stochastic processes occurring at any point up to and including sequencing. One potential explanation of particularly low-abundant taxa detection introduced during sequencing is from index hopping during the sequencing reaction.

**Figure 9.**
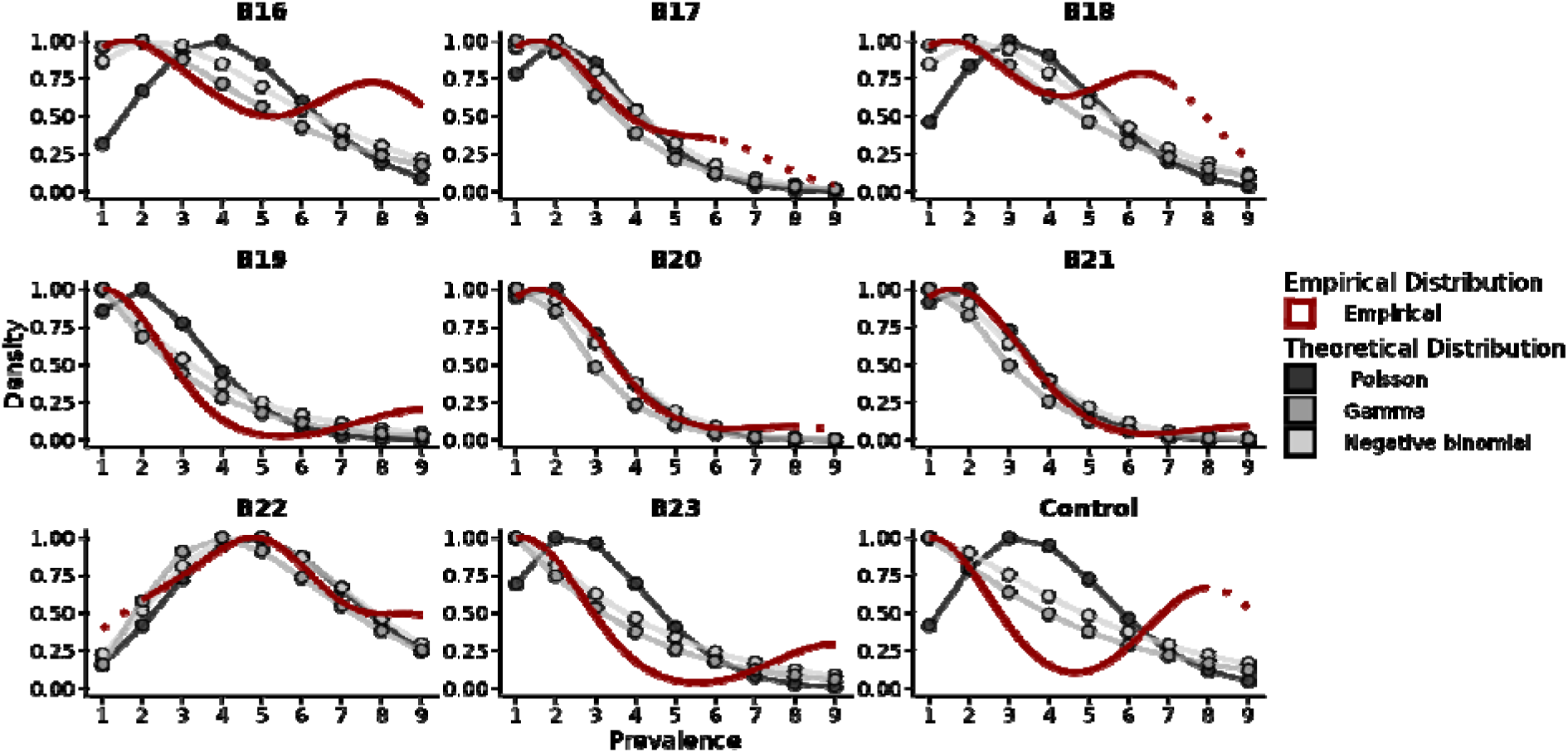
The distribution of within-BALF species prevalence is consistent with a Poisson process. Within-BALF prevalence was calculated for each species and non-zero values plotted to determine the empirical distribution for each BAL sample (red). The theoretical Poisson, gamma, and negative binomial distributions were then fit to the within-BALF species prevalence. The empirical and theoretical distributions of within-BALF prevalence were scaled to a maximum value of 1 to facilitate comparison and interpretation.

### CONSERVED FUNCTIONAL PROFILING SIGNATURES ARE DETECTED ACROSS BALF SAMPLES

In addition to taxonomic profiling, functional profiles were generated at the microbial gene family level (Figure 10A; see Methods for details). Clustering using gene-level Bray-Curtis dissimilarities revealed a densely clustered group of prevalently detected microbial gene families (Figure 10A; top of heatmap). Additional features tended to be more sparsely detected between BALF-samples than features in this cluster. BALF sample B16 had the greatest diversity of sample-specific microbial gene families by visual inspection (Figure 10A). We hypothesize that this densely clustered group of microbial gene families likely stems from highly conserved housekeeping gene families typically found across microbial taxa. Although contaminant microbial gene families were identified and removed using Decontam, it is unclear how efficacious this tool is at removing gene families with reads/signal contributed from both true respiratory tract microbiota and from contaminant taxa. Thus, future efforts to mitigate contamination for functional profiling should likely attempt to remove contaminating sequencing by alignment to metagenome-assembled genomes derived from extraction blanks. In summary, consistent microbial gene family signatures are detected across BALF samples, with some within-BALF sample aliquot heterogeneity; reagent contamination was mitigated by putative contaminant identification, although additional bioinformatic strategies may prove more useful for determining the true functional potential signature in host-depleted lower respiratory tract metagenomes.

**Figure 10.**
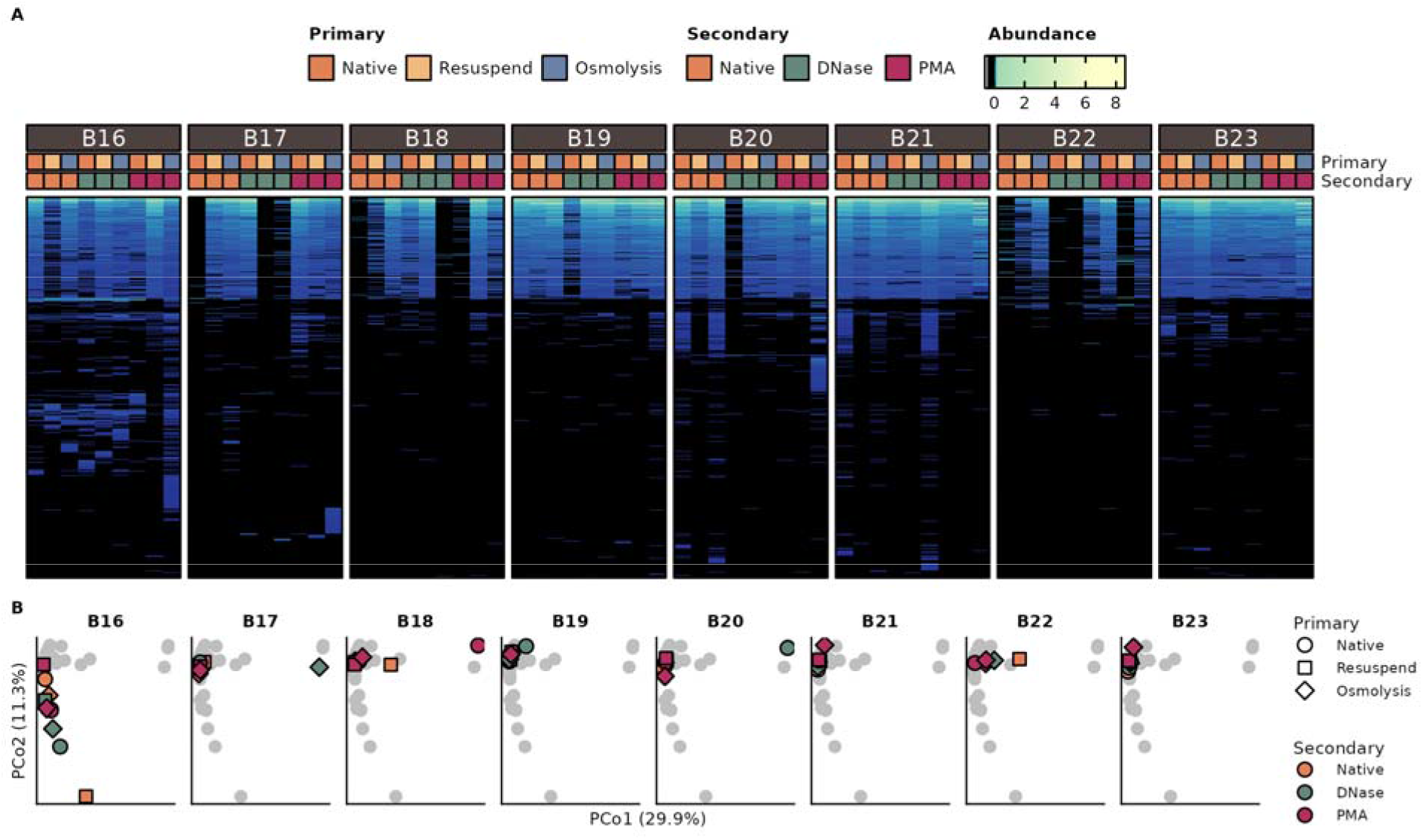
Microbial gene family profiles generated from shotgun metagenomic data. (**A**) Heatmap of functional potential profiles generated using HUMANn3. Rows are hierarchically clustered using Bray-Curtis dissimilarity. Highly abundant microbial gene families clustered at the top of the heatmap are likely derived from highly conserved housekeeping genes. Annotations at the top represent BALF sample and aliquot treatment. Abundances are log_2_ transformed relative abundances with a pseudocount of one. (**B**) Principal coordinate analysis of Bray-Curtis dissimilarities.

Patterns observed in principal coordinate analysis were consistent with the observation of prevalently detected microbial gene families (Figure 10B). That is, samples tended to group closely together in principal coordinate space. Additionally, B16 BALF-sample aliquots had the greatest degree of variation in principal coordinate space (Figure 10B), consistent with the observation that it included the greatest diversity of sparsely detected features (stated above; Figure 10A). Overall, these data indicate that sample to sample heterogeneity tended to be low in functional potential profiles. This is especially notable when compared to principal coordinate analysis of taxonomic profiles, which showed relatively higher sample heterogeneity (Figure 7B).

### DETECTION OF MICROBIAL FEATURES IS ELEVATED WITH PRE-LYSIS TREATMENT COMPARED TO NATIVE CONTROLS

Microbial community profiling yielded variation in feature level detection (Figure 7A and Figure 10A). Data sparsity and inconsistent effects across samples makes overall impact of pre-lysis treatment challenging to interpret. To that end, we compared the average feature abundance in double-native controls to that in treatment aliquots (i.e., NaNa). In taxonomic profiles, we observe that several features were consistently detected at relatively similar abundances between native and treated samples (Figure 11A). While four features were detected in double-native controls that were not detected in processed sample aliquots, many more features were detected in processed samples that were undetectable in native controls (Figure 11A). A similar trend is observed for gene family profiles (Figure 11B). Gene families of lower relative abundances (e.g., less than ∼ 10^-4^) tended to diverge more from the line of perfect association (slope = 1) than gene families detected at greater relative abundances (Figure 11B). Consistent with observations in taxonomic profiles, more features were uniquely detected in processed samples than in double-native controls.

**Figure 11.**
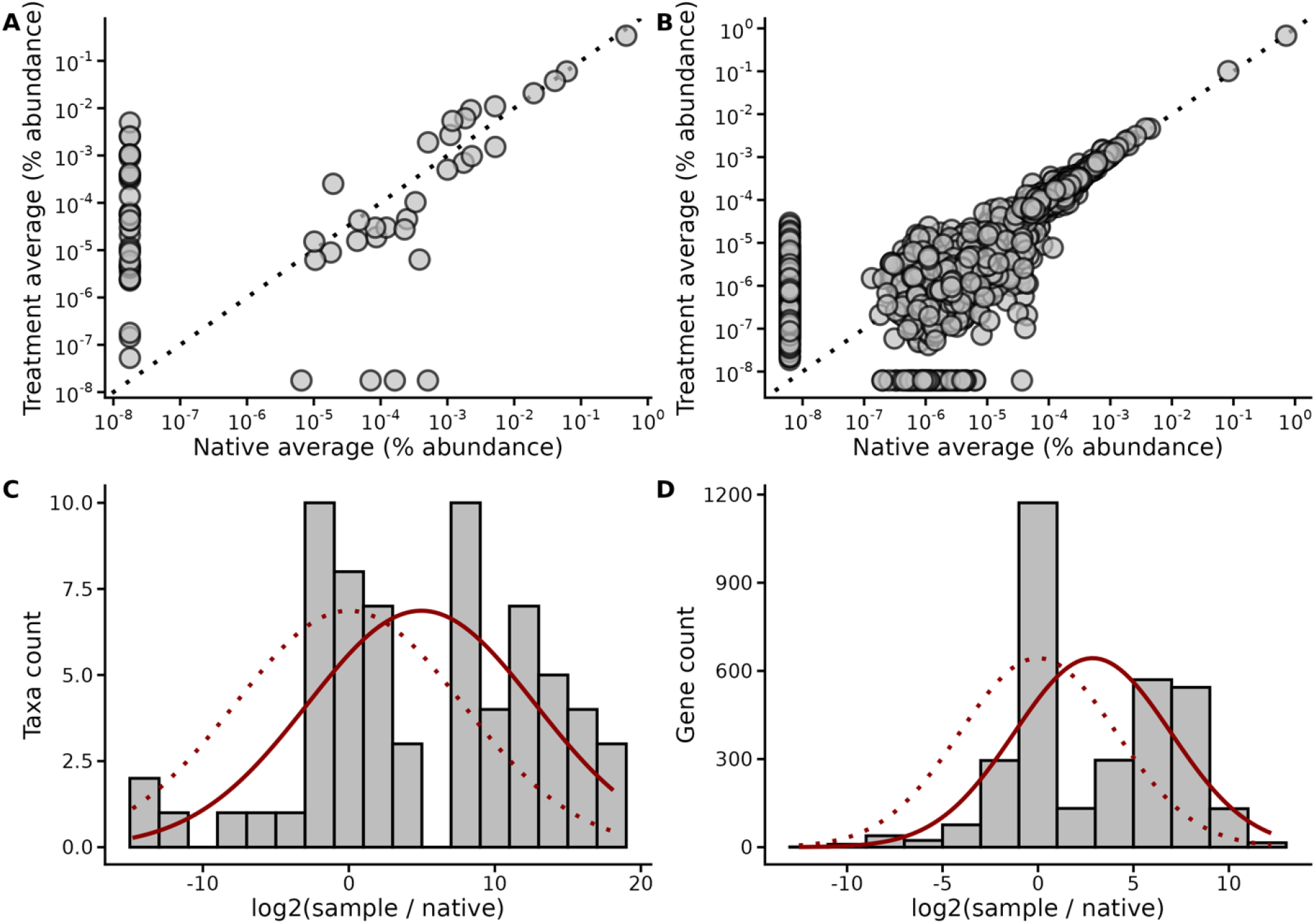
Detection of microbial features is elevated with pre-Lysis treatment compared to native controls. (A-B) Comparison of the mean relative abundance of double native (native-native) treatment controls relative to processed treatment aliquots in (**A**) taxonomic profiles and (**B**) microbial gene famil profiles. Features with a relative abundance of zero were imputed to half the abundance of the lowest detectable abundance in each profile. Dotted lines are indicative of a slope of one between average feature abundance of native and processed sample aliquots. (C-D) Distribution of average feature abundances in treated aliquots relative to native controls in (**C**) taxonomic profiles and (**D**) microbial gene family profiles. Solid red line indicates the fit of the normal distribution to empirical data; dotted lines are the fit of the normal distribution if the mean of the average comparisons were zero. A normal distribution with a mean of zero would be expected by the null hypothesis that microbial community abundance profiles are independent of pre-lysis treatment methodologies. A shifted distribution indicates that community profiles were likely at least partially altered by pre-lysis treatment methodologies.

The number of features detected with respect to relative changes in feature abundance by sample processing was determined for taxonomic (Figure 11C) and microbial gene family profiles (Figure 11D). Under the null hypothesis that feature detection and abundance were independent of treatment processing, one would expect this distribution to fit a normal curve with a mean of zero. However, we observe that this distribution is shifted to the right in both taxonomic and functional profiles. This indicates that community profiles were likely at least partially altered by pre-lysis treatment methodologies.

Additionally, we performed differential abundance analysis on taxonomic profiles using MaAsLin3 (Figure S5). The results showed that the final number of reads after quality control was associated with the greatest number of overabundant features. Several microbial taxa were uniquely depleted in PMA treated samples (e.g., *Klebsiella pneumoniae*; FDR *P* < 0.25), while some others were depleted in in both PMA and DNase treated samples (e.g., *Cutibacterium granulosum*; FDR *P* < 0.25). Of the pre-lysis treatment methods, DNase treatment was the only method to show increased abundance of microbial taxa (i.e., *Enterococcus faecalis, Bifidobacterium breve*; FDR *P* < 0.25). Overall, this analysis indicates that targeting more sequencing depth tended to yield improved microbial coverage and higher microbial abundances.

## 1.4 DISCUSSION

### FRAMEWORK FOR HOST DEPLETION

Host-depletion methods can broadly be categorized into physical or chemical approaches. Physical methods capitalize on conserved differences in cell morphology between mammalian tissue and microbiota. One such characteristic is cell diameter which typically varies greatly between bacteria and their eukaryotic counterparts. Filtration-based cell straining is used to fractionate biological matrixes by particle size, but shear forces, electrostatics, and matrix complexity can reduce separation efficiency. Differential centrifugation applies incremental changes in centrifugal force to pellet particles of different densities. However, this approach is likely to perform poorly because studies have reported a similar and overlapping range of densities between the model bacterium, *E. coli* (∼1.08 g/mL) and human alveolar macrophages (∼1.07 g/mL).^22–24^ Resilience to osmotic pressure is poor in human cells but fair in microbiota due to their prevalent cell walls.^25,26^ Previous studies report that human cells in saliva can be selectively lysed with the addition of water to pelleted samples to improve downstream depletion.^26^ Selective disruption of host cell membranes followed by chemical treatment to deplete freed DNA is a powerful tool for host-depletion.

Chemical depletion methods directly target DNA as opposed to cells themselves. The greatest advantage to this system is the ability to eliminate free DNA for which cell-based targeting fails. Targeted DNA-directed depletion methods discriminate between microbial and host genomes by capitalizing on differentially conserved chemical modifications or sequence homology. Such methods are ideal for samples from which DNA has already been extracted but require further depletion. Methylated cytosines are commonly found at CpG islands in eukaryotic genomes; one commercially available kit (NEBNext Microbiome) targets this modification using paramagnetic-beads linked to MBD2a, a protein that binds methylcytosines, for pulldown-based depletion.^27^ However, they co-target DNA from eukaryotic microbiota with CpG islands. Moreover, 5-methylcytosine modifications are not unique to eukaryotic cells and may also occurs in bacterial genomes thereby limiting selectivity.^28,29^ Homology-based depletion methods using targeted CRISPR-Cas9 cleavage or oligo-mediated pulldown are still in their infancy such that few products are commercially available and rarely target sequences at the genomic scale.^30^ Further advances in homology-based DNA depletion technology is required to mature this promising avenue from gene to genome-scale while retaining rigorous selectivity.

Untargeted DNA-directed depletion methods vary in their capacity to penetrate membranes or matrices. DNase-mediated DNA digestion may readily degrade purified DNA; however, enzymatic activity of wild type proteins like DNaseI suffers severely when used in samples like bronchoalveolar lavage fluid due to, e.g., inhibitor presence. Benzonase and other mutant DNases with resilient profiles have been developed to retain enzymatic activity in unideal conditions.^11,31–34^ Reversible inactivation of Benzonase with EDTA may be considered a risk factor for downstream molecular biology application and irreversible inactivation requires concentrated base at elevated temperatures (100 mM NaOH at 70°C) which is often too extreme for maintaining DNA integrity. Despite this resilience, the sterically bulky nature of enzymes may reduce matrix-penetration and thus depletion efficiency. Propidium monoazide (PMA) is a DNA-binding dye that covalently binds (and potentially cleaves) dsDNA in the presence of light thus sterically hindering polymerases during PCR amplification.^35–40^ Due to its relatively small molecular weight, it readily penetrates disrupted cell membranes and is expected to more readily diffuse across complex matrices. Thus, small-molecule mediated depletion methods may offer a more robust capacity for selective depletion in complex or diffusion-limited systems relative to nuclease digestion.

Enzymatic digestion of cell walls and integral membrane proteins which stabilize cell membranes are classic techniques used for cell lysis. Detergents/surfactants that destabilize lipid bilayers leading to cell lysis usually bias against gram-positives due to their thick peptidoglycan wall. Addition of lysozyme, a peptidoglycan hydrolase, can compensate for this bias yet it does not address the diversity of biopolymers microbes use to build cell walls. A cocktail of lytic enzymes known as MetaPolzyme broadly target diverse cell wall components for degradation to reduce lysis bias at the metagenomic level and yield high molecular weight DNA.^41–43^ While bead-beating is considered a gold standard for metagenomic lysis, limited bias has been previously reported between bead-beating and MetaPolzyme based lysis. Prevalent shearing during bead-beating yields low molecular weight DNA and diminished signal from easily lysed microbes to thus reduce long-read sequencing efficacy and introduce some degradation bias.^41,43^ Therefore, downstream application must be considered when selecting lysis methods due inherent tradeoffs between lysis bias and DNA quality.

### INTERPRETATION OF OUR RESULTS

Screening for effective host depletion using qPCR is a potentially effective strategy for selecting samples likely to be successfully sequenced in metagenomics. In the future, we advise that researchers include ITS2 gene targets for fungal microbiota in addition to 16S rRNA gene and PTGER2 gene targets for their qPCR screens. In several samples, fungal microbiota detection was likely increased by treatment methodology (Figure 2A).

Host depletion gives mixed results that are likely at least partially influenced by sample heterogeneity and the degree of host material. Samples with challenging properties (e.g., presence of red or yellow pigment from biological fluids present in the respiratory tract) tended to be more resistant to improving microbial signal via host depletion (Figure 7A). This indicates that future work should aim to tackle a processing workflow that considers the vast heterogeneity of BALF sample properties from critically-ill patients.

Overall, we observe that chemical depletion, especially treatment with PMA, was able to successfully decrease host-associated biomass (Figure 4F-G). However, this also tended to decrease the bacterial biomass as well, although to a lesser extent than host biomass (Figure 4A-B). Higher levels of bacterial biomass measured using 16S rRNA gene qPCR associated with greater diversity in taxonomic profiles (range FDR *P* < 0.01-0.05; Figure 4C-D). Lower human biomass was independent of microbial profile diversity in our samples. Furthermore, pre-lysis treatment methodologies tended to generate slightly altered microbial community profiles (Figure 6; FDR *P* < 0.05). The differences between these profiles were consistent with the idea that treatment methodologies recover different fractions of the BALF sample microbial community (Figure 6; FDR *P* < 0.05). Overall, this suggests that methodologies to improve sequencing success in BALF samples may find greater success in attempting to enrich microbial DNA load rather than in depleting host DNA load.

Microbial taxa that were inconsistently detected between aliquots of a given BALF sample tended to be of lower relative abundance (Figure 8B). In some cases, certain features were consistently detected at prevalence in one or several BALF samples (Figure 8C-D). Likewise, these taxa were often of low relative abundances compared to taxa with consistent detection signatures (Figure 8E).

To further demonstrate the stochasticity of detecting low-abundant microbial taxa, we draw an analogy to the Poisson process. Typically, a Poisson distribution describes the number of *n* events occurring in a discrete time period if the events occur in manner consistent with a Poisson process (i.e., stochastically).

Here, we model a Poisson distribution using the within-sample prevalence as a discretized analogy to time in a Poisson process. In some instances, species prevalence within a BALF sample was indicative of a Poisson distribution, especially when many taxa were detected at low prevalences (Figure 9). When the empirical distribution for species prevalence within a sample was bimodally distributed, the Poisson distribution was often a poor fit, especially for higher and lower species prevalences. This suggests that in samples with consistent taxonomic signatures, species detection was robust; contrastingly, in samples with sparse taxonomic signatures, rare species detection was likely due to stochastic processes that occurred at some point in the sampling, treatment, and sequencing workflow. However, we cannot exclude the possibility that in some cases a few treatments combination may have been effective at improving microbial profile recovery, which would be expected to fit a Poisson distribution in our analogy to a Poisson process. If this were true, species detected at low prevalence would be expected to be of higher abundance. Although for some features we detect low within-BALF prevalence and moderately high relative abundance, such as *Candida albicans* (Figure 7A), this is not the case for many of the features sporadically detected in our dataset (Figure 8E). Together, this is consistent with the observation that more taxa tended to be detected in processed samples than in double-native controls (Figure 11A-D), although the degree to which they were detected appears spurious. Thus, based on these data, we conclude that low-prevalent and low-abundance taxa were signatures of stochasticity in the experimental workflow. Reducing noise (i.e., background contamination) and improving pre-lysis treatment and processing efficacy may improve the robustness of feature detection in future studies.

Functional profiling yields more consistent detection of a subset of microbial features than does taxonomic profiling (Figure 10). This is consistent with the idea that microbial gene family profiling often yields large amounts of genes with low variation due to being highly conserved across microbiota. Indeed, when modeling the distribution of feature prevalence as a Poisson process, sampling tended to fail to fit a Poisson distribution (Figure S6). This is likely due to multiple microbial taxa contributing to the abundance of a set of shared gene families.

In some cases, host depletion led to contaminants being a greater contributor to the pool of sequenced material (Figure 5). We propose that as host and, to a lesser extent, microbial material is depleted, the relative fraction of reagent contaminants that are co-purified with BALF DNA increases. In addition, variation in contaminant load between reagent lots is likely responsible for the variation in contamination abundance observed in BALF sample aliquots. The former explanation is likely a greater contributor in our dataset, as different BALF samples tended to have similar contamination abundances in no-depletion controls despite performing extractions in randomized batches. In summary, reagent contamination is a critical issue in profiling low-microbial biomass host-associated communities, and host-depletion methodologies tended to promote contamination susceptibility by increasing the fraction of purification space available to contamination.

Based on our results, we present a discussion of what we would recommend to future researchers aiming to improve metagenomic sequencing in BALF samples and other host-associated samples with low microbial biomass. Generally, we recommend that researchers attempt to (1) improve microbial DNA yield, (2) consider all possible avenues for contamination, (3) consider effects of sample heterogeneity.

Liquid-liquid extractions tend to improve bacterial yield in this study and as previously documented.^31,34^ Our data indicate that effectively increasing bacterial yield may generally improve sequencing and microbial profiling success in host-associated microbial communities. Critically, while in our hands, liquid-liquid extraction performed better at bacterial biomass yield (Figure S1), the labor and time costs (which increase potential for introducing contamination) must be considered. An ideal strategy will strive to enhance the yield of microbial biomass while mitigating the potential for contamination through reduced processing time.

We would strongly recommend against the use of DNA-free MetaPolzyme in low-biomass microbial community profiling. Although certified DNA-free, we believe the origin of the majority contaminants in this study (e.g., *Lysobacter enzymogenes*) were derived from this reagent. The DNA-free status of this reagent is determined using PCR amplification with universal 16S and 18S rRNA gene targeting primers, according to the manufacture’s product documentation. Future researchers should be critical of the efficacy of PCR in determining DNA-free certification. More sensitive approaches such as quantitative PCR and high throughput sequencing should be implemented by manufacturers and independently performed by investigative laboratories when using such material in microbiome samples.

BALF samples are notably complex. Individuals tend to harbor different taxonomic signatures.^16^ Visually observable properties of the samples themselves range substantially in (1) viscosity, (2) pigment, and (3) number and size of solid aggregates. We recommend that researchers develop strategies to homogenize BALF sample properties to mitigate the effects of sample properties on host-depletion and/or extraction efficiency. We expect that pigment, especially red pigment derived from blood in the respiratory tract, is likely to reduce photo-activation of PMA. Furthermore, high viscosity fluid may hinder diffusion of DNase or even PMA throughout the sample matrix, reducing overall depletion efficiency. It is unclear at this time whether (1) aggregates carry different biomass yields than non-aggregated BALF and whether (2) microbiota in each fraction is representative of the other. Disruption of aggregates using a sonication bath or exclusion of aggregates altogether may be strategies moving forward depending on the relative yield of human and microbial biomass in BALF aggregates and their overall microbial properties.

## 1.5 CONCLUSION

Effective host depletion in BALF metagenomic sequencing remains an open, critical challenge to the field that will eventually enable cost-effective microbial community reconstruction to inform patient health insights. Methodologies that increase bacterial biomass in BALF samples also tend to resolve microbial communities with greater diversity. We posit that microbial sequencing coverage may be obscured or appear artificially improved by reagent contamination and sequencing artifacts at levels that would be appropriate in many metagenomic sequencing assays but are confounding in samples with low microbial DNA and high human DNA loads. For the evaluation of metagenome sequencing success, it is imperative to assess feature level profiles and read quality control parameters in a transparent manner that is not yet universally adopted across the field. We additionally urge future researchers to consider the potential contamination introduced by biologically derived reagents independent of manufacture certification and the effects of BALF sample heterogeneity on the processing workflow. In summary, we present a framework for improving metagenomic sequencing success in BALF samples and, more generally, in low-biomass, host-associated microbial communities. Host-depletion effects to improve metagenomic sequencing success is a balance between improving microbial signal, mitigating co-purification of reagent contaminants, and canalizing highly heterogenous samples.

## 1.6 ACKNOWLEDGEMENTS

This research was supported in part through the computational resources and staff contributions provided by the Genomics Compute Cluster which is jointly supported by the Feinberg School of Medicine, the Center for Genetic Medicine, and Feinberg’s Department of Biochemistry and Molecular Genetics, the Office of the Provost, the Office for Research, and Northwestern Information Technology. The Genomics Compute Cluster is part of Quest, Northwestern University’s high performance computing facility, with the purpose to advance research in genomics.

## 1.7 DATA AND CODE AVAILABILITY

Sequencing data were deposited onto SRA BioProject PRJNA1366628. Code is available on the GitHub repository: <u>hartmann-lab/Sumner2025_BALF_Methods</u>.

## 1.8 METHODS

### SAMPLE ACQUISITION AND PROCESSING

#### Sample Acquisition

Patient samples were collected as part of the Successful Clinical Response to Pneumonia Therapy study which focuses on mechanically ventilated patients in the intensive care unit; this study received approval the Northwestern University Institutional Review Board (STU00204868). Bronchoalveolar lavage was performed on critically ill patients with pneumonia by appropriate health care providers at Northwestern Medicine. Procedures were performed using standard of care practices. Excess bronchoalveolar lavage fluid not used for clinical testing (∼10 mL) was stored at 4°C in 15 mL falcon tubes until downstream processing, typically within 1 day post-procedure. Samples were homogenized by inversion and gentle vortexing before aliquoting out into at least 10 aliquots of 500 µL using autoclaved wide-bore tips. After aliquoting bronchoalveolar lavage fluid, samples were stored on ice while not being actively processed. All treatment processing steps, with the exception of heat treatments and centrifugation, were performed in a biosafety cabinet until lysis for DNA extraction. Biosafety cabinet and micropipettes were surface decontaminated using 70% ethanol followed by UV-irradiation. Regular exchange of gloves was implemented as previously suggested for processing BALF samples.

#### Primary Treatment Processing

Primary treatment was performed on fresh BALF sample aliquots. Three native controls were set aside on ice. Concurrently, 6 treatment aliquots were centrifuged at 5,000xg for 10 minutes at 4°C in a benchtop centrifuge. A subset of 3 pelleted aliquots were resuspended by percussive agitation of the tube (flicking) and returned to ice. The approximately 450 µL of supernatant for the remaining 3 aliquots was carefully decanted using a p200 micropipette and discarded. Pellets were resuspended with 450 µL of nuclease free water by pipetting and briefly vortexing, followed by incubation at room temperature for 5 minutes.

#### Secondary Treatment Processing

Secondary treatment was performed on aliquots from each of the primary treatment processing groups (i.e., combinatorially). Secondary treatment groups also included a native control, where aliquots were set on ice and processed no further. The main treatment groups were either incubation with PMA or DNaseI.

For PMA treatment, 5 µL of 2 mM PMA solution (Biotium, catalog no. 40019) was deposited onto the lid of each tube in a dark room, for a final concentration of 20 µM PMA. After addition of PMA, tubes were transferred to a freezer box covered in aluminum foil and inverted approximately 50 times to mix PMA droplet with BALF sample. Aliquots were set on a ThermoMixer shaking at 300 RPM for 10 minutes before. Aliquots were transferred to a PMA-Lite device and exposed to light for PMA activation for 15 minutes. Every 2 minutes, samples were removed, briefly vortexed, and returned to the PMA-lite device; device and timer were momentarily paused during vortexing to consistently monitor light exposure. Samples were then wrapped in foil before storage at −80°C.

For DNase treatment, 5 µL of DNase I was added to each aliquot and incubated at 37°C on a heat block for 20 minutes. A subsequent 75°C incubation for 10 minutes was used to inactivate DNaseI activity. The time for the heat block to reach the inactivation temperature, approximately 6 minutes, was not counted against heat inactivation incubation time. Sample aliquots were then cooled on ice before storage at −80°C.

### DNA PURIFICATION

#### Setup and Batch Randomization for Liquid-Liquid Extraction

DNA was purified from processed BALF sample aliquots using a liquid-liquid extraction methodology based on the Lucigen MasterPure Complete DNA and RNA Purification kit (catalog no. MC85200).^34^ Extractions were performed in 7 batches of up to eleven aliquots and at least one negative extraction control; sample were assigned to each batch in a randomized manner to reduce the potential for batch effect biases. First, BALF sample aliquots were thawed on ice. During this time, three 1.5 mL microcentrifuge tubes corresponding the (1) lysate, (2) intermediate precipitation, and (3) purified DNA rehydration were prepared and labelled for each purification sample before opening BALF sample aliquots to reduce potential for cross-contamination from aerosols or the environment. After BALF sample aliquots thawed, samples were homogenized by vortexing and collecting residual volume by brief centrifugation in a picofuge. Then, 250 µL of each BALF sample aliquots was transferred to a fresh microcentrifuge tube (“lysate”). Additionally, 250 µL DPBS was transferred to a fresh tube to be used as a blank extraction control. Details of the remaining two tubes are described in the paragraphs directly below.

#### Bacterial Cell Lysis with MetaPolzyme

For each extraction batch, one vial of certified DNA-free MetaPolzyme (Sigma-Aldrich, catalog no. MAC4LDF) was resuspended in 100 µL of sterile DPBS by gentle pipetting and set aside in 4°C until ready for enzymatic digestion. Residual MetaPolzyme was stored at 4°C when not fully consumed and prioritized for use in the next batch. To each sample aliquot, 10 µL of MetaPolzyme was added and homogenized by slowly inverting approximately six times followed by a brief vortex. Enzymatic digestion was performed on a ThermoMixer at 37°C rotating at 500 RPM for 75 minutes.

#### RNA and Protein Digestion

After bacterial cell lysis, samples were cooled on ice for approximately 5 minutes. During this time, a master mix of PK/TC solution was generated by diluting 1.66 µL of proteinase K into 250 µL 2x T and C Lysis Solution. Condensate and residual lysate were collected from the sides of sample tubes by briefly centrifuging sample aliquots. Then, 250 µL of PK/TC solution was added to each sample followed by briefly vortexing. Proteinase K digestions was performed at 65°C for 25 minutes with rotation at 300 RPM. All samples were cooled to 37°C in the ThermoMixer (approximately 10 minute ramp down) followed by addition of 1.66 µL of RNase A (5 mg/mL) and incubation for 50 minutes, still shaking at 300 RPM. Samples were placed on ice for approximately 5 minutes to cool before continuing to DNA precipitation.

#### DNA Precipitation

Samples were vortexed and briefly centrifuged to collect residual liquid; then, 300 µL lysate was transferred to a fresh tube. The remaining lysate was frozen at −80°C. Protein Precipitation Reagent was added at a volume of 150 µL to each sample and vigorously vortexed in pulses for 10 seconds. Precipitate was pelleted from solution by centrifugation with 16,000xg at 4°C for 18 minutes, with additional centrifugation performed as needed when it was evident that precipitate had not fully sedimented/was still in solution. Approximately 420 µL supernatant was transferred to a new microcentrifuge tube (“purified DNA”) and the pellet was discarded. A 1x volume of isopropanol was added to recovered supernatant followed by briefly vortexing and 50-60x inversions in a freezer box (Fisherbrand™ Cryo/Freezer Boxes, catalog no. 03-395-464). DNA was then precipitated by centrifugation with 16,000xg at 4°C for 18 minutes and the supernatant was subsequently decanted into the discard. Next, an ethanol wash was performed twice using fresh 70% ethanol; ethanol was dispensed on the side of the tube opposite of the pellet to mitigate disruption. Between washes, samples centrifuged for 2 minutes at 16,000xg and removal of the supernatant. Residual ethanol was removed using a p20 micropipette prior to vacuum centrifugation for 2 minutes with V-AQ mode to desiccate. DNA was resuspended in 50 µL of nuclease free water and stored at 4°C to rehydrate and eventually to storage at −80°C. DNA concentration was assessed using the fluorometric Quant-iT dsDNA High Sensitivity Assay Kit (Invitrogen, catalog no. Q33120).

### QUANTITATIVE PCR ASSAYS

#### Cloning Standards for qPCR

A fragment of the PTGER2 gene was amplified from human DNA (Roche) and cloned into pCR4-TOPO vector using the TOPO-TA kit according to the manufacturer instructions. The PTGER2 gene was amplified using the previously published primers (F: 5’-GCTGCTTCTCATTGTCTCGG-3’ and R: 5’-GCCAGGAGAATGAGGTGGTC-3’).^26^ NEB 5-alpha competent cells were transformed and plated on tryptic soy agar plates with kanamycin (50 µg/mL) selection. For ten colonies, one-half colony was picked using an autoclaved pipette tip and inoculated into 20 µL of PCR master mix (10 µL 2x KAPA HiFi, 1 µL of PTGER2 Primer Mix [8 µM of forward and of reverse primer], and 9 µL of UltraPure water) to screen for insertion of the expected target sequence. The remaining half of each colony was inoculated into Luria-Bertani broth. Plasmids were extracted from overnight cultures derived from colonies with positive-screen results using the QIAprep Spin Miniprep Kit (Qiagen, catalog no. 27104). Glycerol stocks of *E. coli* with vector inserts of the 16S rRNA gene were previously prepared^34^ and plated under selection, grown in liquid culture, and miniprepped as described above. Cloned sequences were confirmed using nanopore whole plasmid sequencing (Primordium labs).

#### qPCR Methods

Universal primers targeting the bacterial 16S rRNA gene (341F: 5’-CCTACGGGAGGCAGCAG-3’ and 818R: 5’-ATTACCGCGGCTGCTGG-3’)^44^ and the human PTGER2 gene (described above) were used to amplify conserved genomic regions in qPCR assays and thus serve as a proxy for genomic copies. Absolute quantification was performed using the standard curve method in both qPCR-based assays. Standard curves were constructed ranging from 10^8^ to 10^2^ copies of plasmid standard per qPCR reaction. Reactions were comprised of 10 µL 2x PowerUp HotStart SYBR Mix (BioRad, catalog no. A25780), 1 µL of Primer Mix (8 µM of forward and of reverse primer), and 9 µL of template diluted 1:9 in UltraPure water. Assays, including standard curve and sample dilution, were performed using an EpMotion5073m liquid handling robot. A QuantStudio3 qPCR thermocycler was used for reaction.

### SHOTGUN METAGENOMIC SEQUENCING ASSAY

#### Library Construction, Pooling, and Sequencing Shotgun Sequencing

Metagenomic sequencing was used to assess effects of host-depletion on microbial community profiling. In addition to BALF samples and extraction blanks, a no-template control (i.e., nuclease-free water) and a ZymoBIOMICS Microbial Community DNA Standard (catalog no. D6305) were included as negative and positive controls for library construction. Shotgun metagenomic libraries were prepared using NEBNext® Ultra™II FS DNA Library Prep Kit for Illumina (NEB catalog no. E7805L) following manufacturers’ instructions. Each sample library was constructed using 500pg input DNA template or up to 10 µL of material if the concentration was too low. Adapters were added using the IDT® for Illumina® DNA/RNA UD Indexes Set A, Tagmentation (catalog no. 20027213) for tagmentation. Library quality and quantity was assessed by Tapestation High Sensitivity D1000 (Agilent Technologies) and Qubit dsDNA HS Assay Kit (Invitrogen, catalog no. Q32851). Libraries were pooled in an equimolar ratio for multiplexed sequencing. All available material was included for pooling when libraries were of insufficient quantity. Pooled libraries were sequenced on a NextSeq2000 instrument with a P3 flow cell (catalog no. 20027800) for a sequencing reaction of 2×151 cycles with 8×8 cycles for adapter sequencing.

### BIOINFORMATICS

#### Sequencing Quality Control

Sequencing data intensities were deconvolved into FASTQ files using BCL convert (v.4.3.6). Read quality control was then performed using KneadData (v.0.12.0), which implements several quality control tools. First, adapter sequences and low-quality bases were removed using Trimmomatic (v.0.33); an adapter sequence file was supplied that included standard adapter sequences including the universal Illumina adapters, TruSeq3 adapters, and NexteraPE adapters.^45^ Additionally, a 20-mer repeating G sequence was included to target possible polyG tails typical for NextSeq2000 instruments and the ‘--run-trim-repetitiv’ flag to remove overrepresented sequences identified in the initial FastQC screening. Reads less than 30bp after adapter trimming were excluded. Execution of Tandem Repeat Finder (TRF) was bypassed using the ‘--bypass-trf’ flag. Finally, reads were serially aligned the human reference genome (CHM13 Telomere-to-Telomere with Y chromosome from GRCh38) and a ribosomal gene sequence database (SILVA 128 LSUParc SSUParc ribosomal RNA database) using Bowtie2 (v.2.5.4).^46^ The number of read passing each step were tracked and visualized in Figure 3B.

#### Taxonomic & Functional Profiling

Short read taxonomic profiling was performed with MetaPhLAn4 (v.4.1.1).^47^ Clean reads were processed using MetaPhLAn4 with the vJan25 ChocoPhlAn database; default parameters were used except for a minimum read length of 45bp and a marker gene abundance percentile threshold of 0.2 (‘--stat_q’). Taxonomic profiles were analyzed at the single genome bin (SGB) level, the highest taxonomic resolution available in MetaPhLAn4.

Functional abundance profiles were then generated using HUMAnN3 (v.3.9).^48^ As the vJan25 database was not compatible with HUMAnN3 at the time of analysis, a second set of MetaPhLAn4 profiles were generated using the older vJun23 ChocoPhlAn database for pan-genome nucleotide database construction. Reads were mapped to the pan-genome HUMAnN3 ChocoPhlAn database (v.201901_v31) using Bowtie2 nucleotide alignment; unmapped reads were then processed using the UniRef90 database with translated search, requiring at least half the subject sequence to be covered (‘–translated-subject-coverage-threshold 50.0’). Microbial gene pathway abundances were computed using default parameters. Microbial gene family abundances were normalized to relative abundance. Microbial gene families, which default to UniRef90 terms, were regrouped into PFAM domain. Normalized PFAM abundance profiles were then unstratified (i.e., using the overall microbial gene family abundance rather than the species-specific microbial gene family abundance) for downstream processing.

#### Putative Contaminant Identification

Putative contaminants were identified using the Decontam R package (v.1.26.0).^49^ Briefly, the prevalence method was used to compare samples to negative extraction controls. The positive ZymoBIOMIC Microbial Standard sample and the library construction no-template control were excluded from Decontam processing as contaminants were expected to be introduced during DNA extraction. All negative extraction controls were included for contaminant identification. The threshold parameter was set to 0.1 for contaminant selection. Additional species of the genus *Lysobacter* and the species *Methylibium petroleiphilum* were manually included as contaminants. The human microbiota of *Staphylococcus pasteuri* and the uncultured Lawsonellaceae member (SGB3663) were excluded from the contaminant list as they were of consistent low abundance among blank extractions and are potentially derived from BALF samples. Decontamination of HUMAnN3 functional potential profiles was also performed using the prevalence method with a threshold of 0.1 in the Decontam R package.

#### Analysis of Alpha and Beta Diversity

Alpha and Beta diversity calculations were performed using various functions from the Vegan R package (v.2.7-1). Alpha diversity was assessed for taxonomic profiles using three metrics: Shannon, inverse Simpson, and richness. The Shannon index and the inverse Simpson index were calculated using the *diversity* function. Richness calculation was performed using the *specnumber* function from the same package. Sample beta diversity was determined using the *vegdist* function; both Bray-Curtis dissimilarities and Jaccard distances were computed. Principal coordinate analysis was then performed with the *cmdscale* function from the stats R package (v.4.4.0) on beta diversity metrics; eigenvalues explaining the amount of variance by each principal coordinate were extracted for plot annotation with the *eigenvals* function from Vegan.

### STATISTICAL ANALYSES

#### Data Visualization & Frequency Tests

Statistical analyses were performed in R (v.4.4.0). Data were primarily visualized using the ggplot2 R package (v.3.5.2). Additional visualization utilities used were from the ggpubr (v.0.6.1), ggh4x (v.0.3.1), and patchwork (v.1.3.1) R packages. Unless otherwise stated, boxplots were configured such that the center line is representative of the median value, box limits represent the upper and lower quartiles, and whiskers are 1.5x the interquartile range.

Several analyses involved comparison of quantitative data between categorical groups (e.g., 16S rRNA gene copy number between treatments in Figure 3). As previously stated in their respective figure caption, statistical significance was assessed for these comparisons using the non-parametric Wilcoxon Signed-Rank test for paired data and the Wilcoxon Rank-Sum test for unpaired data as implemented in the rstatix R package (v.0.7.2). Multiplicity correction was performed using the Benjamini-Hochberg method.

#### Multivariate Analysis

Permutational analysis of variance (PERMANOVA) was performed to determine associations between treatment conditions and other metadata with microbial community composition and functional potential. PERMANOVA analysis was performed with the *adonis2* function; a total of 4,999 permutations were executed for each test. Permutations were restricted within individual BALF samples but freely allowed permutations between aliquots, unless otherwise stated. Multiplicity correction was performed using the Benjamini-Hochberg method for each dataset. Co-variation between different profiles was assessed using the Mantel test as implemented in the ade4 R package (v.1.7-23) using 5,000 permutations.

#### Differential Abundance Analysis

Differential abundance analysis was performed using MaAsLin3 (v.0.99.18).^50^ Secondary treatment, primary treatment, and the number of cleaned high-quality reads were included as fixed effects. BALF sample index was used as a random effect value and replaced as a fixed effects terms when the number of observations for a given feature was small. Unscaled absolute abundances from 16S rRNA gene qPCR assays were for modeling absolute abundance variation. As 16S gene copies is specific to bacterial absolute abundance, eukaryotic taxonomic features were excluded from differential abundance testing. Features with low prevalence (0.05) and low variance (0.1) were also excluded from testing. For normalization and transformation settings, the default TSS parameter and the non-default pseudo-log parameter with a zero-threshold of negative one were selected. An alpha of 0.25 was used to determine statistical significance.

## 1.9 SUPPLEMENTAL TEXT

### PRELIMINARY LYSIS SCREEN, METHODOLOGY

After collection the 8 BALF samples and before proceeding with the extraction methodology described in the main text, we set out to identify an optimal method for improving bacterial load yield. We hypothesized that improved bacterial recovery, independent of host depletion, would be an important factor for lower respiratory tract communities metagenomic sequencing. Our experimental approach and results from qPCR screening is described in the paragraphs directly below.

#### Pooling Samples for Typical Representation

From each of the eight BALF samples, one untreated aliquot of 500 µL was thawed one ice and used for generating a representative BALF samples. BALF aliquots were triturated (pipetted up and down) to break apart mucosal aggregates and pooled into a 5 mL conical tube. The conical tube was inverted and gently vortexed to homogenize the sample prior to distributing in 250 µL aliquots across twelve 1.5 mL microcentrifuge tubes. Samples were stored on ice while not being processed.

#### Preliminary Lysis and Extraction Methodology

MetaPolzyme, lysozyme, bead-beating, and detergent lysis was performed on the generated representative BALF sample aliquots in triplicate. In addition to each treatment group, blank extractions were performed in parallel to the detergent extraction, using 250 µL nuclease-free water instead of representative BALF. To prepare MetaPolzyme enzyme solution, vials of certified DNA-free MetaPolzyme (Sigma) were resuspended in 100 µL of Dulbecco’s Phosphate-Buffered Saline (DPBS) with gentle pipetting. To prepare the lysozyme solution, 1g of lysozyme (Sigma, catalog no. L6876) was suspended in 100 µL of 10 mM Tris-HCl, pH 8.0 for a final concentration of 10 mg/mL lysozyme. Resuspended enzyme solutions were stored at 4°C until ready for use. For lysis with enzymatic reagent, 10 µL of enzyme solution (either MetaPolzyme or lysozyme) was added to each aliquot and set on ice prior to incubation. Sample aliquots in the detergent-only group were supplemented with 10 µL of DPBS and set on ice prior to incubation. MetaPolzyme, lysozyme, and detergent-only aliquots were then incubated at 37°C at 500 RPM for 60 minutes on a ThermoMixer.

The bead-beating procedure was performed in a similar manner as previously described.^34^ A lysis buffer solution was generated by supplementing 250 µL of 2X T and C Lysis Solution (Lucigen) with 1.66 µL of proteinase K (PK/TC solution). Representative BALF aliquots were transferred into an O-ring tube with varying sized beads (Zymo) with 250 µL of PK/TC solution. Bead-beating was performed in a BioSpec Mini-Beadbeater-96 (catalog no. 1001) for 5 minutes. Samples were stored on ice while remaining samples were incubated as described in the above paragraph.

A magnetic bead-based approach was also assessed and automated for use with BALF sample DNA extraction. The Zymo Quick-DNA™ HMW MagBead Kit (catalog no. D6060) was selected for DNA extraction because of its potential for using in long read sequencing. Extractions were performed according to the manufacturer’s instructions. After enzymatic digestions were set on ice, 250 µL BTSB buffer solution and 20 µL proteinase K were added to each aliquot tube and incubated at room temperature for 20 minutes. Subsequently, samples were briefly centrifuged to collect liquid and transferred to a 96 deep-well plate for remaining extraction steps. Remaining steps of the extraction procedure were adapted for automation in the EpMotion5073M liquid handling device.

After incubation was complete for enzymatic lysis and detergent treatments, 250 µL of PK/TC solution was added to each tube. Then, all 12 samples were incubated at 60°C for 25 minutes at 300 RPM on a ThermoMixer, vortexing samples every 5 minutes. Lysed samples were centrifuged at 12,000xg for 5 min, namely, to collect beads from bead-beating treatment, and 300 µL lysates were transferred to a new microcentrifuge tube. RNaseA was then added in a 3.33 µL volume to each lysate followed by vortexing and a 37°C incubation for 50 minutes at 300 RPM. Following RNaseA incubation, samples were set on ice for 5-8 minutes. Then, 300 µL of each sample was transferred to a new tube. Protein Precipitation Reagent was added at a volume of 0.5x to each aliquot followed by vigorous vortexing for 10 seconds. Precipitate was pelleted from solution by centrifugation with 16,000xg at 4°C for 18 minutes. Supernatant was recovered into a fresh microcentrifuge tube at approximately 420 µL and nucleic acids were precipitated with using approximately 1x volume of chilled isopropanol. Centrifugation was performed as for protein precipitation. Supernatant was then carefully decanted and an ethanol wash was performed twice using fresh 70% ethanol and centrifugation for 2 minutes at 16,000xg. Residual ethanol was removed using a p20 micropipette prior to vacuum centrifugation for 3 minutes with V-AQ mode to desiccate. DNA was resuspended in 50 µL of nuclease free water and incubated at 4°C overnight to rehydrate.

#### Preliminary qPCR Standard Curve Generation

*E. coli* strain ATCC 25922 was revived from a lab glycerol stock by plating on Tryptic Soy Agar followed by incubation at 37°C. Colonies were selected for inoculation of Tryptic Soy Broth and overnight growths were performed at the same temperature under shaking conditions. Genomic DNA was extracted using the GenElute™ Bacterial Genomic DNA Kit (Millipore Sigma, cat no. NA2100) as per the manufacturer’s guidelines. DNA concentration was quantified using the Quant-iT assay. A standard curve of known 16S rRNA gene copies ranging from 1×10^8^ to 1×10^2^ copies per reaction was generated using genomic *E. coli* DNA by assuming a genome size of 5.2 Mb and a 16S rRNA gene copy number of 7 *E. coli* per genome. Quantitation of the 16S rRNA gene was otherwise performed as described below.

### SCREENING LYSIS METHODOLOGY FOR BACTERIAL BIOMASS YIELD, RESULTS

We first sought to identify factors that would improve overall bacterial biomass yield in metagenomic DNA extraction from BALF samples. We identified four distinct lysis treatment groups that we hypothesized would show differential yield on bacterial biomass: (1) physical lysis with bead-beating, (2) enzymatic lysis with MetaPolzyme, (3) enzymatic lysis with lysozyme, and (4) chaotropic lysis with detergent alone. Due to the rarity of BALF samples and the extent of our experimental plans, eight BALF samples were combined in equal volumes to generate a representative BALF mixture. To test our hypothesis of differential bacterial yield, aliquots of the representative sample were generated, and subsequently processed for each lysis group in triplicate.

Furthermore, we expected extraction methodology to impact overall DNA yield. To that end, we attempted to test the various lysis methods using a high-labor liquid-liquid DNA extraction strategy (Lucigen MasterPURE) and an automated magnetic bead-based extraction strategy (Zymo MagBead). All lysis treatment groups were based on the same extraction kit, and the extraction procedure converged after the group-specific lysis strategy. To that, each group was subject to incubation with lysis buffer followed by proteinase-K treatment in addition to their physical or enzymatic lysis strategy.

16S rRNA gene copy concentration was largely similar between the lysis-treatment groups (FigS1). The 16S rRNA gene was quantified at approximately 1.5×10^5^ copies in each reaction for the Lucigen-extracted samples, independent of the lysis methodology. The Zymo-extracted samples yielded approximately less than 2.5×10^4^ 16S rRNA gene copies in each reaction across lysis methodologies. The observation that Lucigen MasterPure extractions improve quantity of bacterial DNA isolation is consistent with previous reports.^31,34^ The lysis group with the greatest bacterial load, although only slightly, was MetaPolzyme-lysis with Lucigen extraction. For this reason, and the potential to pursue long-read sequencing with high molecular weight DNA, remaining samples were extracted using this strategy with slight alterations.

### FILTERING WAS UNSUCCESSFULLY ATTEMPTED AS A PRIMARY TREATMENT METHOD

We hypothesized that host cells could be separated from bacterial cells on account of the relative difference in cell size using a sterile mesh filter. To that end, we attempted filtration in the following manner. Sterile PluriSelect cell strainers were fitted to a 50 mL falcon tubes with connector ring attachments. After addition of 1.5 mL of bronchoalveolar lavage fluid, 1.0 mL syringes were inserted into the connector ring attachment to promote straining under low pressure. As bronchoalveolar lavage fluid often retains viscous clumps, likely originating from biofilms or mucosal aggregates, most samples failed to strain. For this reason, we excluded filtration from downstream analysis.

## 1.10 SUPPLEMENTAL FIGURES

**Supplemental Figure 1.**
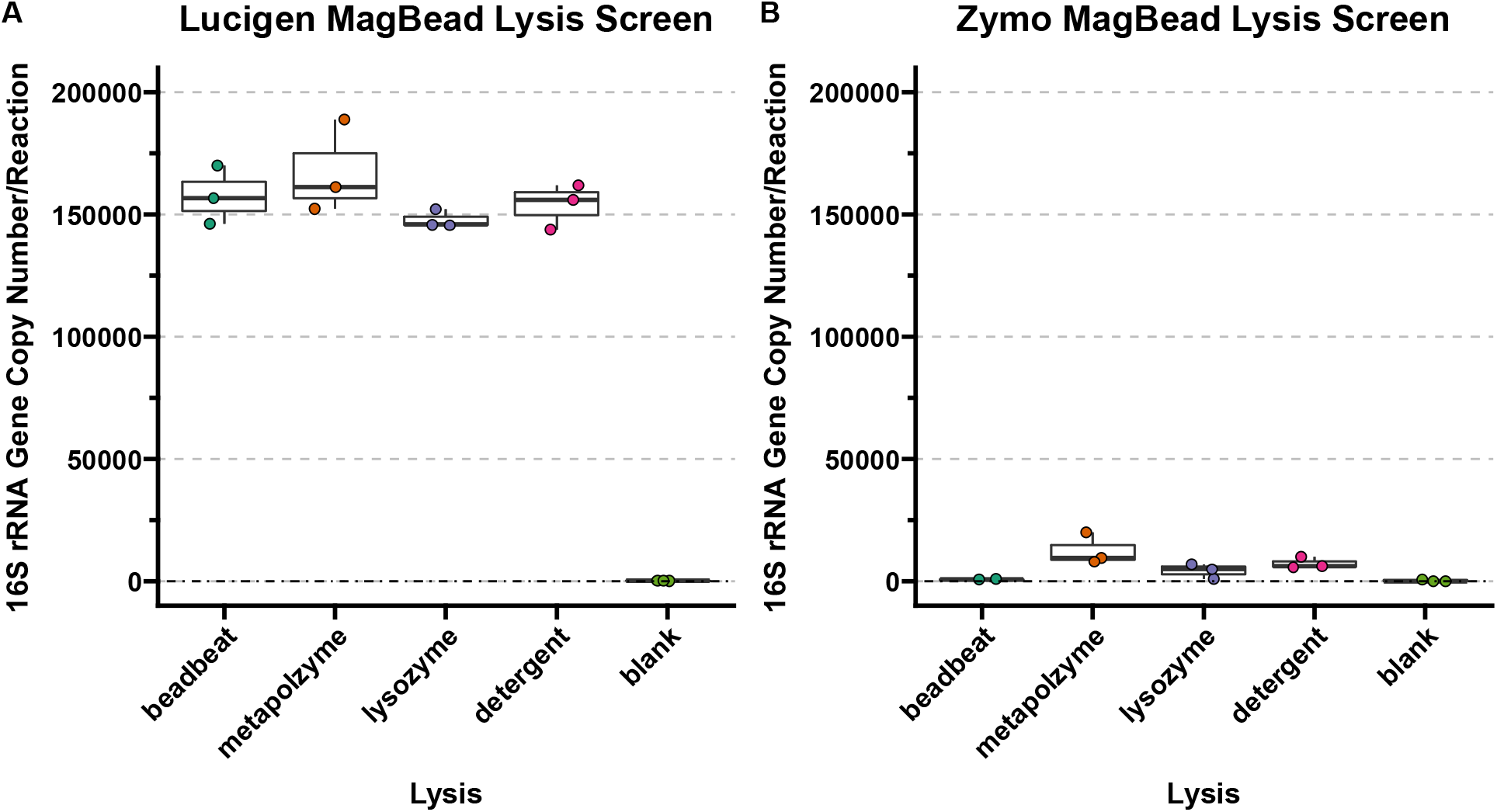
Impact of lysis method on bacterial biomass yield. Lysis-screen experiment performed using (**A**) Lucigen MasterPURE (manual) and (**B**) Zymo MagBead Extraction kits (automated). Equal volumes of BAL samples were thawed, pooled, and aliquoted to create a representative BAL. In technical triplicate, aliquoted samples were subjected to primary lysis using either enzymatic (MetaPolzyme, lysozyme), physical (bead-beating), or shams. Following primary lysis, detergent-based lysis was performed with proteinase k digestion. Blank (i.e., water) controls were used to control for environmental microbiota.

**Supplemental Figure 2.**
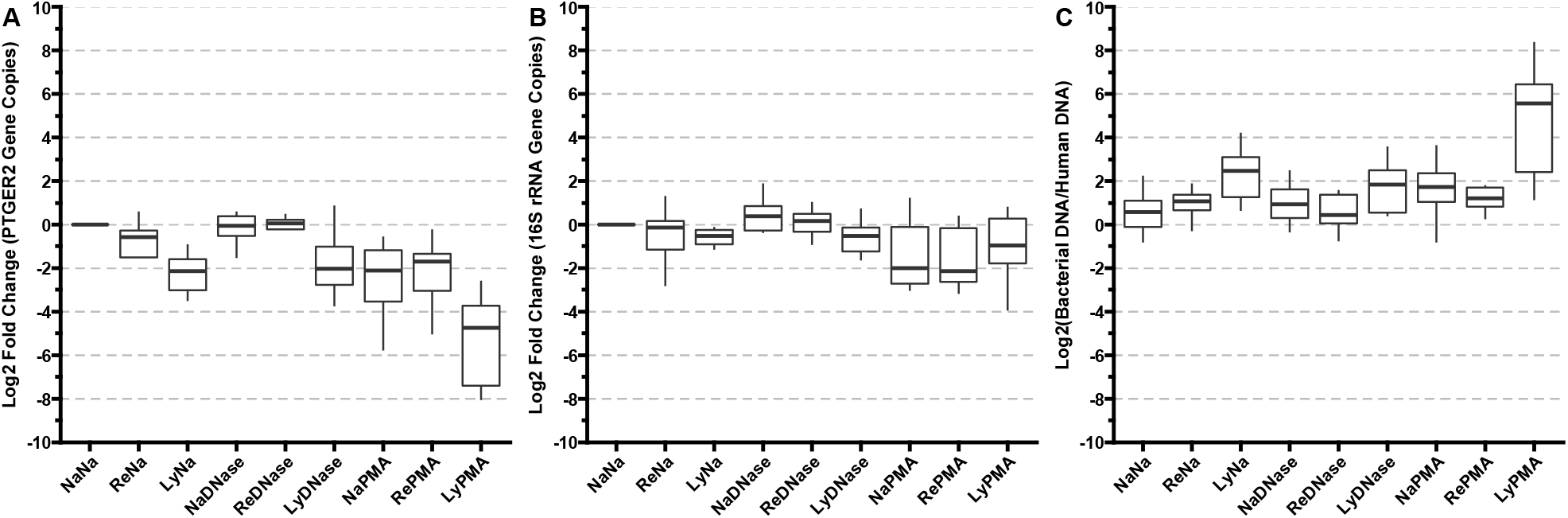
Quantitative PCR analysis of treatment effect on patient and microbial DNA yield. (A) Treatment effect on PTGER2 copy number relative to baseline control. (B) Treatment effect on 16S rRNA gene copy number relative to baseline control. (C) Ratio of 16S rRNA gene to PTGER2 gene copy number shows the normalized proportion of estimated bacterial DNA to human DNA.

**Supplemental Figure 3.**
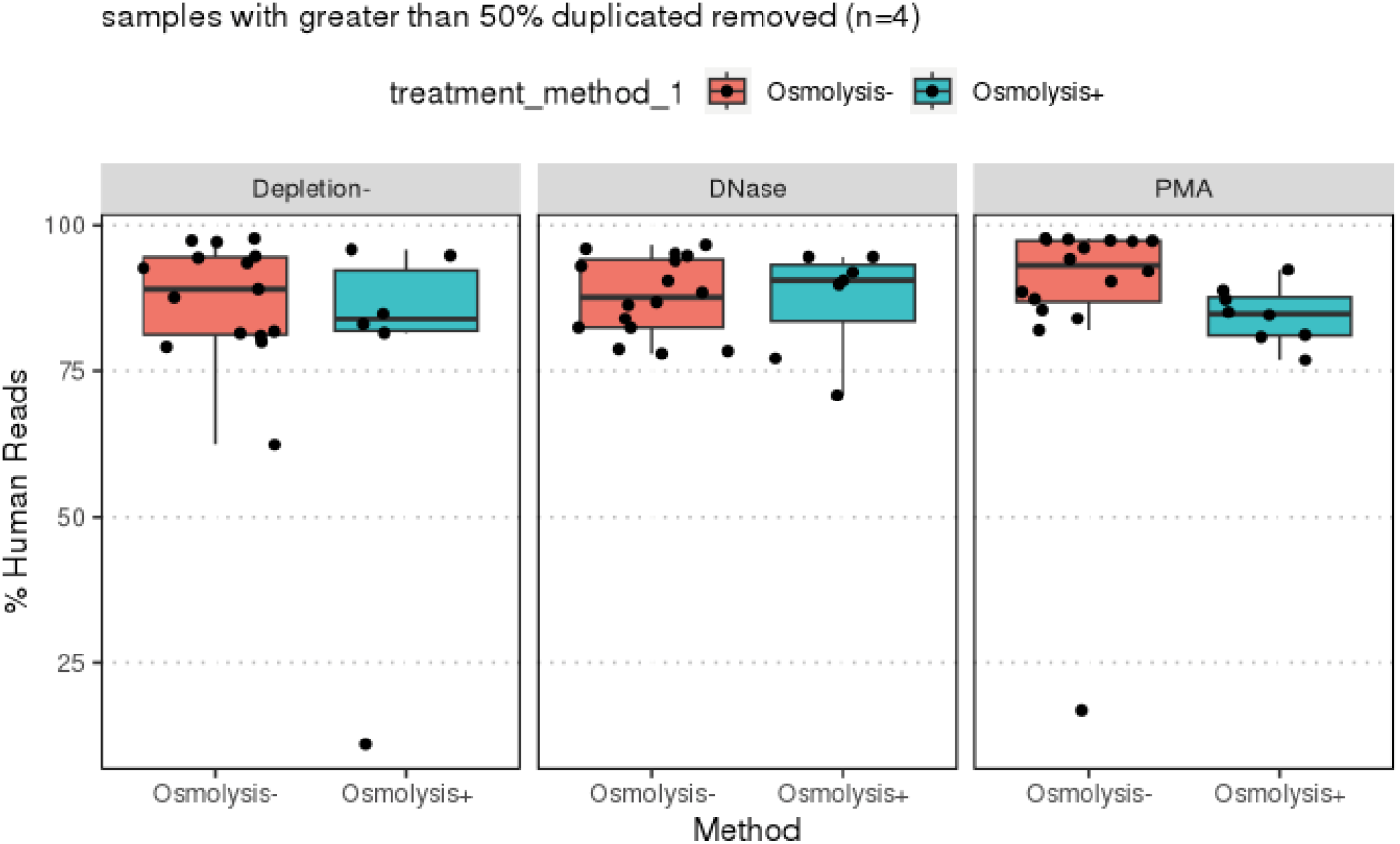
Shotgun sequencing analysis of treatment effect on patient and microbial DNA yield. Variation in the fraction of human reads assayed from metagenomic sequencing libraries was observed.

**Supplemental Figure 4.**
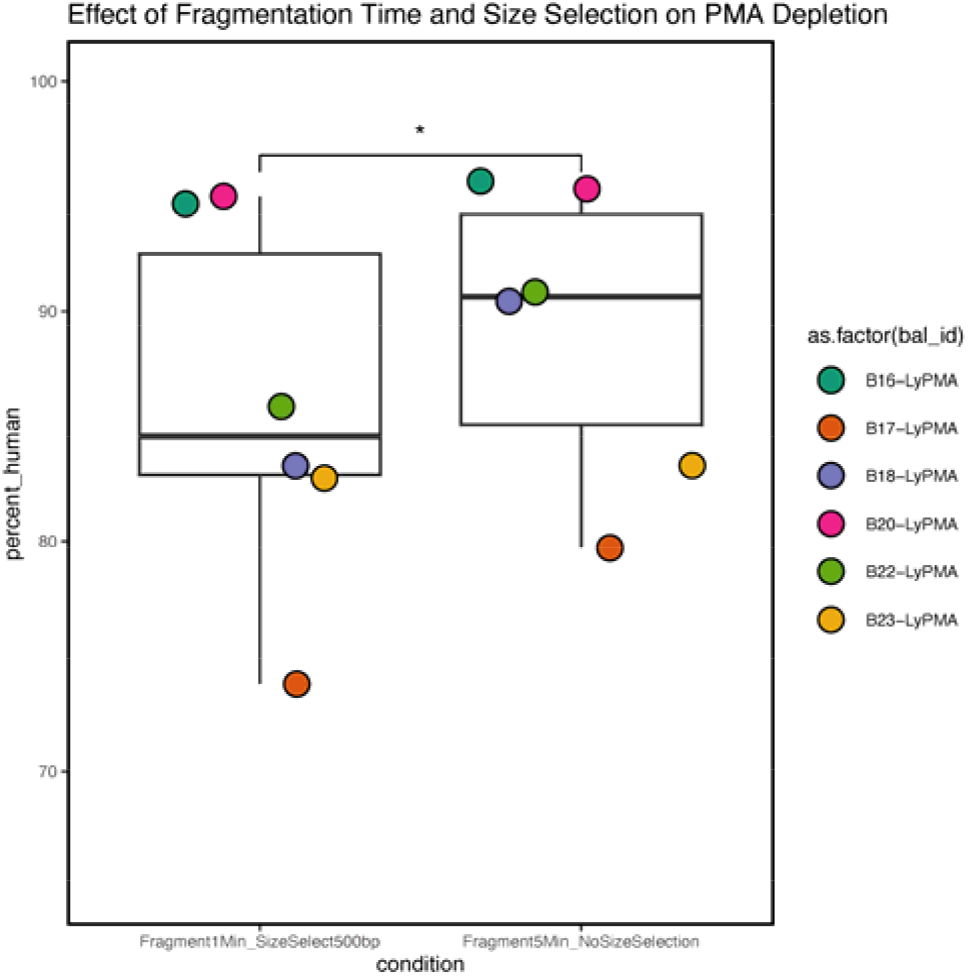
Effect of library preparation fragmentation on fraction of human reads in metagenomic sequencing data. Reads were processed using fastp and aligned to the human reference genome with Bowtie2.

**Supplemental Figure 5.**
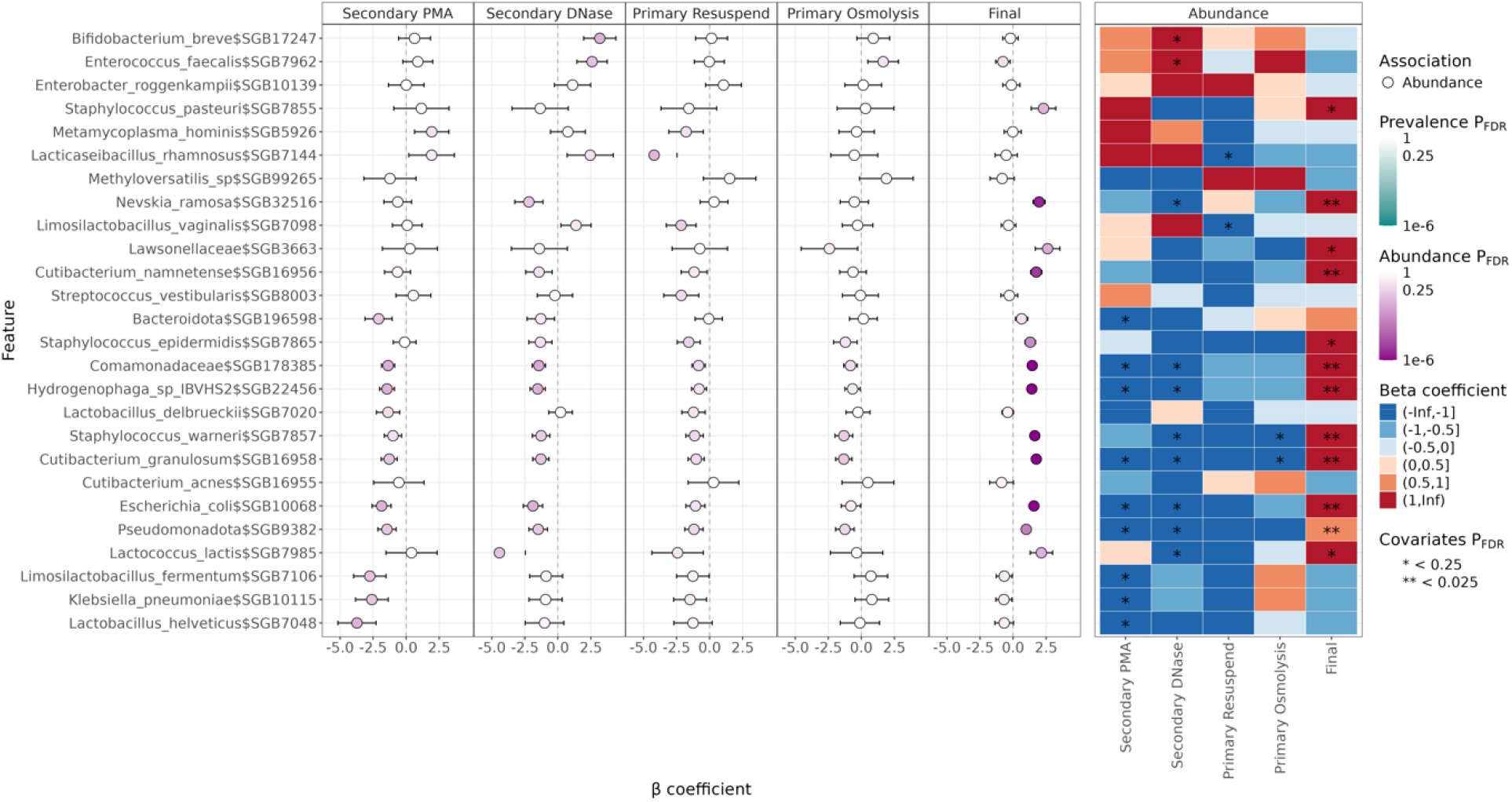
Results of differential abundance testing using MaAsLin3. Differential abundance analysis was performed using absolute abundance estimation on the bacterial fraction of species-level MetaPhlAn4 profiles. The final read count after quality control with KneadData is indicated by “Final.”

**Supplemental Figure 6.**
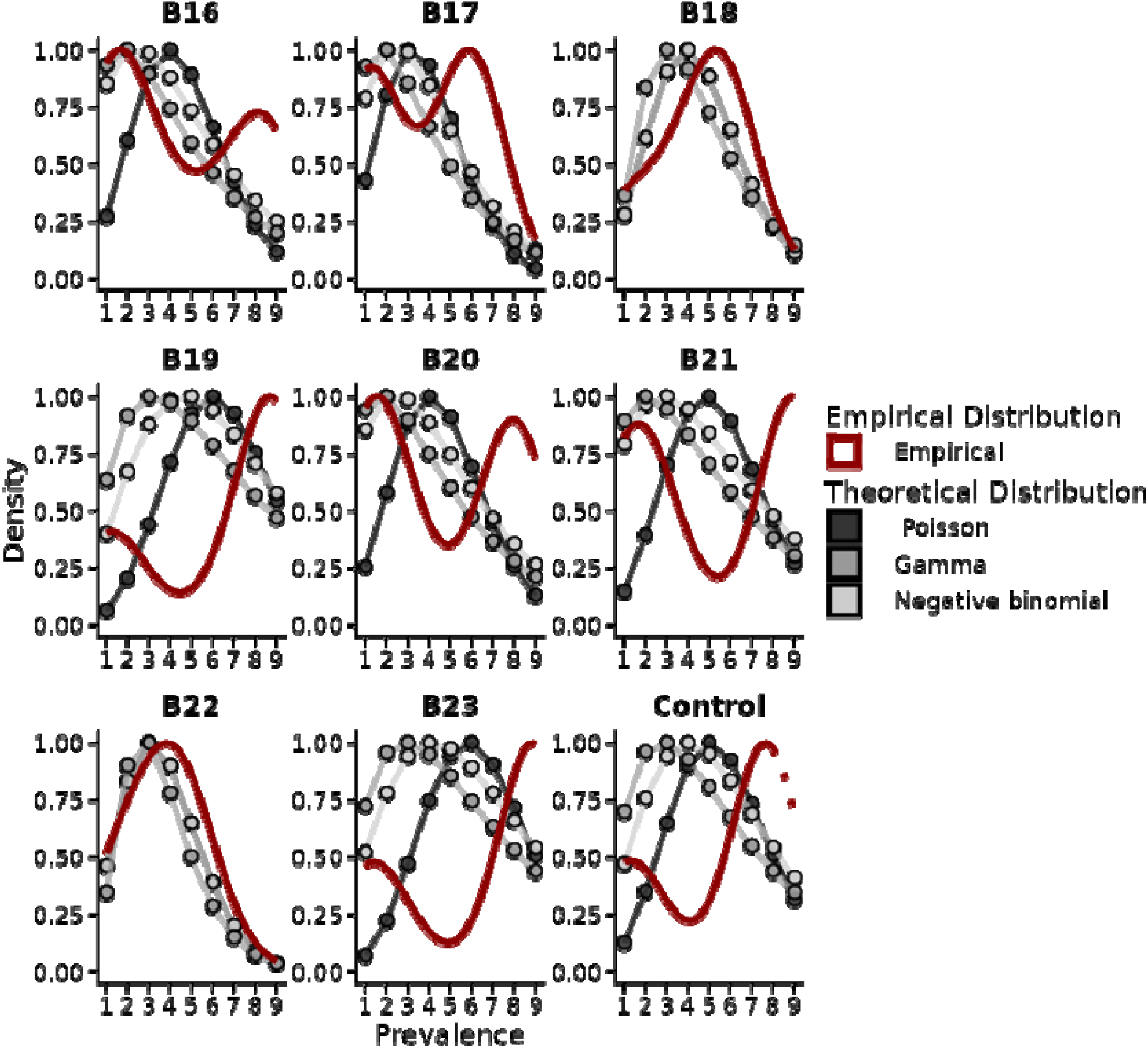
The distribution of within-BALF microbial gene family prevalence is inconsistent with a Poisson process. Within-BALF prevalence was calculated for each microbial gene family and non-zero values plotted to determine the empirical distribution for each BAL sample (red). The theoretical Poisson, gamma, and negative binomial distributions were then fit to the within-BALF gene famil prevalence. The empirical and theoretical distributions of within-BALF prevalence were scaled to a maximum value of 1 to facilitate comparison and interpretation.

## 1.11 SUPPLEMENTAL TABLES

**Supplemental Table 7.** Sequencing quality control statistics.

**Supplemental Table 2.** Meta’omic sequencing profiles from MetaPhlAn and HUMAnN.

**Supplemental Table 3.** Contaminant identification results.

**Supplemental Table 4.** Quantitative PCR results.

